# Updating the sulcal landscape of the human lateral parieto-occipital junction provides anatomical, functional, and cognitive insights

**DOI:** 10.1101/2023.06.08.544284

**Authors:** Ethan H. Willbrand, Yi-Heng Tsai, Thomas Gagnant, Kevin S. Weiner

## Abstract

Recent work has uncovered relationships between evolutionarily new small and shallow cerebral indentations, or sulci, and human behavior. Yet, this relationship remains unexplored in the lateral parietal cortex (LPC) and the lateral parieto-occipital junction (LPOJ). After defining thousands of sulci in a young adult cohort, we revised the previous LPC/LPOJ sulcal landscape to include four previously overlooked, small, shallow, and variable sulci. One of these sulci (ventral supralateral occipital sulcus, slocs-v) is present in nearly every hemisphere and is morphologically, architecturally, and functionally dissociable from neighboring sulci. A data-driven, model-based approach, relating sulcal depth to behavior, further revealed that the morphology of only a subset of LPC/LPOJ sulci, including the slocs-v, is related to performance on a spatial orientation task. Our findings build on classic neuroanatomical theories and identify new neuroanatomical targets for future “precision imaging” studies exploring the relationship among brain structure, brain function, and cognitive abilities in individual participants.

## Introduction

A fundamental goal in psychology and neuroscience is to understand the complex relationship between brain structure and brain function, as well as how that relationship provides a scaffold for efficient cognition and behavior. Of all the neuroanatomical features to target, recent work shows that morphological features of the shallower, later developing, hominoid-specific indentations of the cerebral cortex (also known as putative tertiary sulci, PTS) are not only functionally and cognitively meaningful, but also are particularly impacted by multiple brain-related disorders and aging (Amiez et al., 2019, 2018; Ammons et al., 2021; Cachia et al., 2021; Fornito et al., 2004; Garrison et al., 2015; Harper et al., 2022; Lopez-Persem et al., 2019; Maboudian et al., 2024; Miller et al., 2021; Nakamura et al., 2020; Parker et al., 2023; Ramos Benitez et al., 2024; Voorhies et al., 2021; Weiner, 2019; Willbrand et al., 2023b, 2022a, 2022b; Yao et al., 2022). The combination of these findings provides growing support for a classic theory proposing that the late gestational emergence of these PTS in gestation within association cortices, as well as their prolonged development, may co-occur with specific functional and microstructural features that could support specific cognitive abilities that also have a protracted development (Sanides, 1964). Nevertheless, despite the developmental, evolutionary, functional, cognitive, and theoretical relevance of these findings, PTS have mainly been restricted to only a subset of association cortices such as the prefrontal, cingulate, and ventral occipitotemporal cortices (Amiez et al., 2019, 2018; Ammons et al., 2021; Cachia et al., 2021; Fornito et al., 2004; Garrison et al., 2015; Harper et al., 2022; Hathaway et al., 2023; Lopez-Persem et al., 2019; Miller et al., 2021, 2020; Nakamura et al., 2020; Parker et al., 2023; Voorhies et al., 2021; Weiner, 2019; Willbrand et al., 2023b, 2023c, 2022a, 2022b; Yao et al., 2022). Thus, examining the relationship among these PTS relative to architectonic and functional features of the cerebral cortex, as well as relative to cognition, remains uncharted in other association cortices such as the lateral parietal cortex (LPC).

As LPC is a cortical extent that has expanded extensively throughout evolution (Van Essen et al., 2018; Zilles et al., 2013), there is great interest in the structure and function of LPC in development, aging, across species, and in different patient populations. Yet, key gaps in knowledge relating individual differences in the structure of LPC to individual differences in the functional organization of LPC and cognitive performance remain for at least four main reasons. First, one line of recent work shows that LPC displays a much more complex sulcal patterning than previously thought (Drudik et al., 2023; Petrides, 2019; Segal and Petrides, 2012; Zlatkina and Petrides, 2014), while a second line of work shows that LPC is tiled with many maps and discrete functional regions spanning modalities and functions such as vision, memory, attention, action, haptics, and multisensory integration in addition to theory of mind, cognitive control, and subdivisions of the default mode network (Goodale and Milner, 1992; Harvey et al., 2015, 2013; Humphreys and Tibon, 2023; Konen and Kastner, 2008; Mackey et al., 2017; Schurz et al., 2017). Second, a majority of the time, the two lines of work are conducted independently from one another and the majority of human neuroimaging studies of LPC implement group analyses on average brain templates—which causes LPC sulci to disappear (**Fig. 1**). Third, despite the recently identified complexity of LPC sulcal patterning, recent studies have also uncovered previously overlooked PTS in association cortices (for example, in the posterior cingulate cortex; Willbrand et al., 2023c, 2022a). Thus, fourth, it is unknown if additional LPC PTS are waiting to be detailed and if so, could improve our understanding of the structural-functional organization of LPC with potential cognitive insights as in other association cortices. Critically, while such findings would have developmental, evolutionary, functional, cognitive, and theoretical implications for addressing novel questions in future studies, they would also have translational applications as sulci serve as biomarkers in neurodevelopmental disorders (Ammons et al., 2021; Cachia et al., 2021; Garrison et al., 2015; Nakamura et al., 2020) and “corridors” for neurosurgery (Tomaiuolo and Giordano, 2016).

**Fig. 1.**
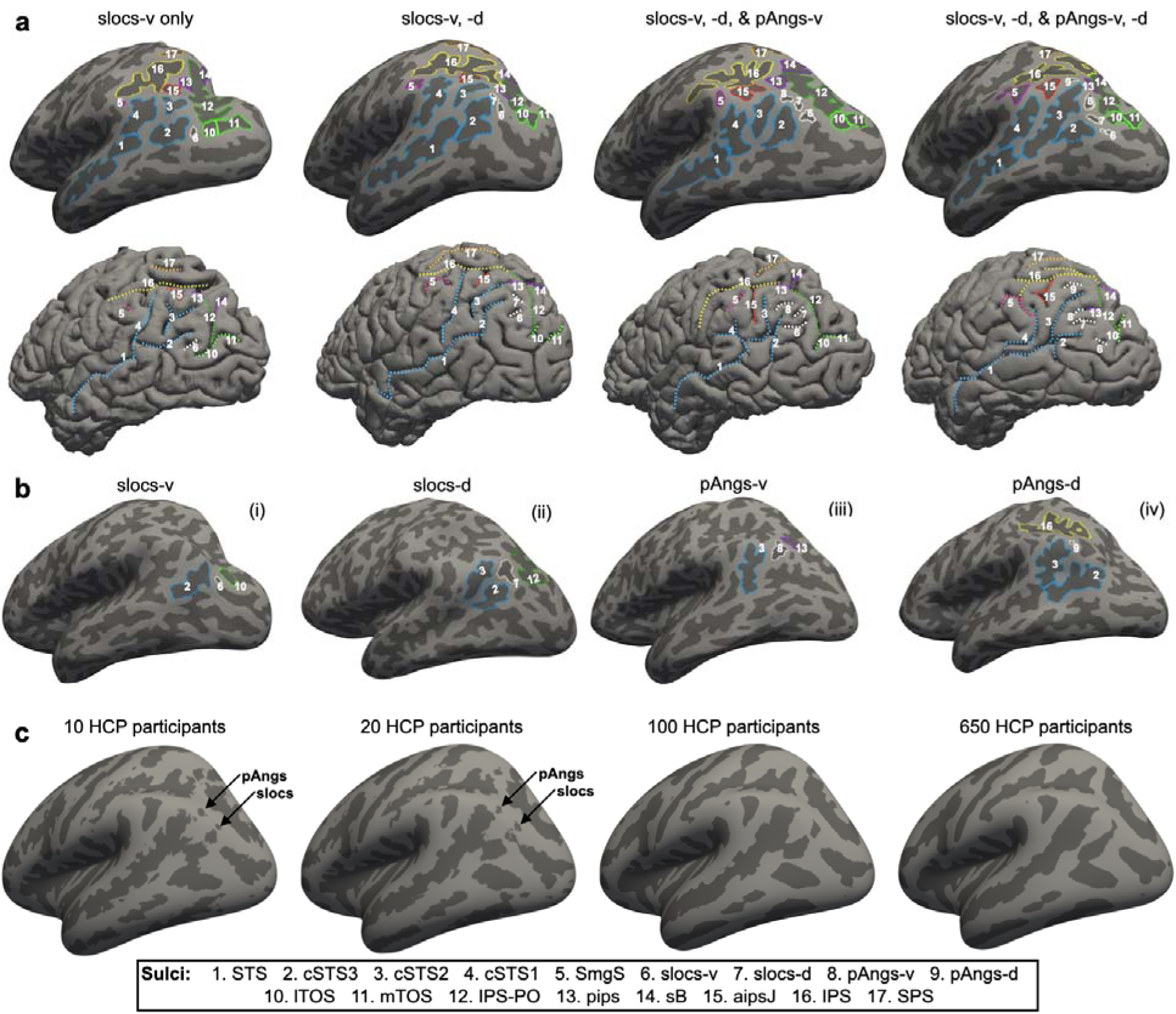
Four previously undefined small and shallow sulci in the lateral parieto-occipital junction (LPOJ). **a.** Four example inflated (top) and pial (bottom) left hemisphere cortical surfaces displaying the 13-17 sulci manually identified in the present study. Each hemisphere contains 1–4 of the previousl undefined and variable LOC/LPOJ sulci (slocs and pAngs). Each sulcus is numbered according to the legend. **b.** Criteria for defining slocs and pAngs components. (i) Slocs-v is the cortical indentation between the cSTS3 and lTOS. (ii) Slocs-d is the indentation between cSTS3/cSTS2 and IPS-PO. (iii) pAngs-v is the indentation between the cSTS2 and pips. (iv) pAngs-d is the indentation between cSTS2/cSTS1 and IPS. **c.** The variability of the slocs and pAng components can cause them to disappear when individual surfaces are averaged together. Left to right: (i) 10 Human Connectome Project (HCP) participants, (ii) 20 HCP participants, (iii) 100 HCP participants, and iv) 650 HCP participants. The disappearance of these sulci on average surfaces, which are often used for group analyses in neuroimaging research, emphasizes the importance of defining these structures in individual hemispheres.

In the present study, we first manually defined LPC sulci in 144 young adult hemispheres using the most recent definitions of LPC sulci (Petrides, 2019). By manually labeling over 2,000 sulci, we detail four previously undescribed (Supplementary Methods and Supplementary Figs. 1–4 for historical details) sulci in the cortical expanse between the caudal branches of the superior temporal sulcus (cSTS) and two parts of the intraparietal sulcus (IPS)—a cortical expanse recently referenced as containing sensory “bridge” regions of the temporal-parietal-occipital junction (Glasser et al., 2016)—which we term the supralateral occipital sulci (ventral: slocs-v; dorsal: slocs-d) and posterior angular sulci (ventral: pAngs-d; dorsal: pAngs-d). We then utilized morphological (depth and surface area), architectural (gray matter thickness and myelination), and functional (resting-state functional connectivity) data available in each participant to assess whether the most common of these structures (slocs-v) was dissociable from surrounding sulci. Finally, we assessed whether the updated view of the LPC/LPOJ sulcal landscape provided cognitive insights using a model-based, data-driven approach (Voorhies et al., 2021) relating sulcal morphology to behavior on tasks known to activate regions within this cortical expanse (for example, reasoning and spatial orientation; Gur et al., 2000; Karnath, 1997; Vendetti and Bunge, 2014; Wendelken, 2014).

## Results

### Four previously undescribed small and shallow sulci in the lateral parieto-occipital junction (LPOJ)

In previous research in small sample sizes, neuroanatomists noticed shallow sulci in this cortical expanse, but did not describe them beyond including an unlabeled sulcus in their figures and did not consider individual differences (Supplementary Methods and Supplementary Figs. 1–4 for historical details). In the present study, we fully update this sulcal landscape considering these overlooked indentations. In addition to defining the 13 sulci previously described within the LPC/LPOJ, as well as the posterior superior temporal cortex in individual participants (**Methods**; Petrides, 2019), we could also identify as many as four small and shallow PTS situated within the LPC/LPOJ that were highly variable across individuals and left undescribed until now (Supplementary Methods and Supplementary Figs. 1–4). Though we officially name and characterize features of these sulci in this paper for the first time, it is necessary to note that the location of these four sulci is consistent with the presence of variable “accessory sulci” in this cortical expanse mentioned in prior modern and classic studies (Supplementary Methods). For four example hemispheres with these 13-17 sulci identified, see **Fig. 1a** (Supplementary Fig. 5 for all hemispheres).

Macroanatomically, we could identify two sulci between the cSTS3 and the IPS-PO/lTOS ventrally and two sulci between the cSTS2 and the pips/IPS dorsally. Ventrally, we refer to these sulci as ventral (slocs-v; sulcus 6 in **Fig. 1**) and dorsal (slocs-d; sulcus 7 in **Fig. 1**) components of the supralateral occipital sulcus (slocs). The slocs-v, located between the posterior cSTS3 and lTOS, was present in 98.6% of hemispheres (left hemisphere: N = 71/72; right hemisphere: N = 71/72; **Fig. 1**). Conversely, the more variable slocs-d, located between the cSTS3 and IPS-PO, was present 68.0% of the time (left hemisphere: N = 50/72; right hemisphere: N = 48/72; **Fig. 1**). Dorsally, we refer to the other newly described sulci as the ventral (pAngs-v; sulcus 8 in **Fig. 1**) and dorsal (pAngs-d; sulcus 9 in **Fig. 1**) components of the posterior angular sulcus (pAngs). The pAngs components were more rare than the slocs components. Specifically, pAngs-v, located between cSTS2 and pips, was identifiable 31.3% of the time (19 left and 26 right hemispheres; **Fig. 1**). Located between cSTS2 and the IPS, pAngs-d was identifiable only 13.2% of the time (8 left and 11 right hemispheres; **Fig. 1**). These incidence rates were significantly different (GLM, main effect of sulcus: χ2(3)L=L166.53, *p* < .0001; no hemispheric effects: *p*s > .68). The slocs-v was more common than the other three sulci (*p*s < .0001), slocs-d was more common than the pAngs components (*p*s < .0001), and pAngs-v was more common than pAngs-d (*p* = .002). We could also identify these sulci in post-mortem hemispheres (Supplementary Figs. 2, 3), ensuring that these sulci were not an artifact of the cortical reconstruction process.

Beyond characterizing the incidence of sulci, it is also common in the neuroanatomical literature to qualitatively characterize sulci on the basis of fractionation and intersection with surrounding sulci (termed “sulcal types”; for examples in other cortical expanses, see Chiavaras and Petrides, 2000; Drudik et al., 2023; Miller et al., 2021; Paus et al., 1996; Weiner et al., 2014; Willbrand et al., 2022a). All four sulci most commonly did not intersect with other sulci (see **Supplementary Tables 1–4** for a summary of the sulcal types of the slocs and pAngs dorsal and ventral components). The sulcal types were also highly comparable between hemispheres (rs > .99, *p*s < .001).

Given that sulcal incidence and patterning is also sometimes related to demographic features (Cachia et al., 2021; Leonard et al., 2009; Wei et al., 2017), subsequent GLMs relating the incidence and patterning of the three more variable sulci (slocs-d, pAngs-v, and pAngs-d) to demographic features (age and gender) revealed no associations for any sulcus (*p*s > .05). Finally, to help guide future research on these newly- and previously-classified LPC/LPOJ sulci, we generated probabilistic maps of each of these 17 sulci and share them with the field with the publication of this paper (**Fig. 2**; **Data availability**).

**Fig. 2.**
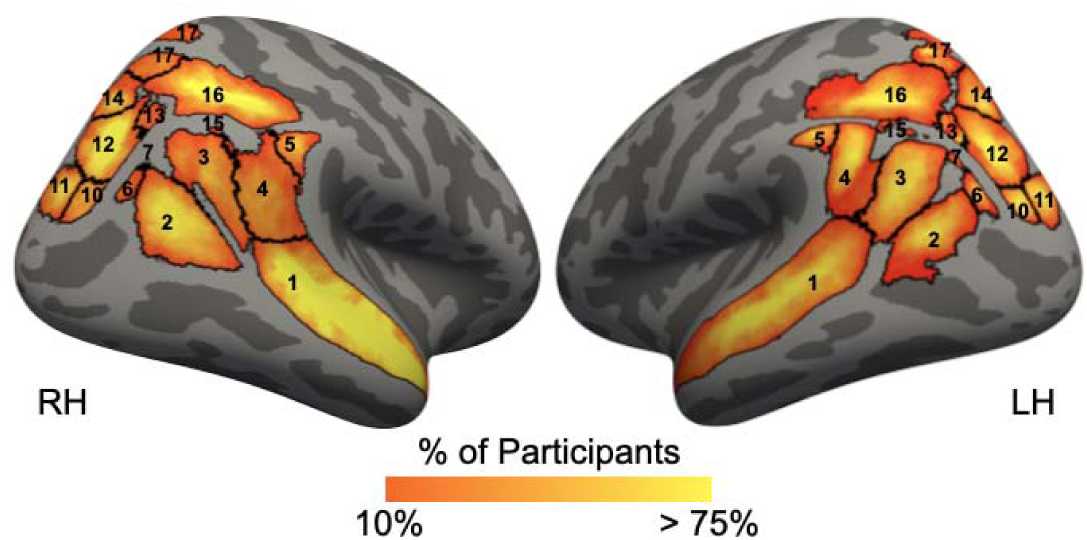
Maximum probability maps of the 17 sulci identified in the present study. Maximum probability map (MPMs) for the 17 LPC/LPOJ sulci on the inflated fsaverage cortical surface (sulci: dark gray; gyri: light gray; cortical surfaces are not to scale) in the left (right surface; LH) and right (left surface; RH) hemispheres. To generate the MPMs, each label was transformed from each individual to the fsaverage surface. For each vertex, the proportion of participants for whom that vertex i labeled as the given sulcus (the warmer the color, the higher the overlap) was calculated. In the cases in which the vertices for each component overlapped, the sulcus with the highest overlap across participants was assigned to that vertex. For visual clarity, the MPMs were thresholded to 20% overlap acros participants. Sulci are numbered according to Fig. 1. These sulcal MPMs can be used to guide the definition of LPC/LPOJ sulci in future studies.

### The slocs-v is morphologically, architecturally, and functionally dissociable from nearby sulci

Given that the slocs-v was present in the majority of participants (98.6% across hemispheres) and the other three sulci were far more variable (<70% of hemispheres), we focused our analyses on this stable sulcal feature of the LPOJ. To do so, we first tested whether the slocs-v was morphologically (depth and surface area) and architecturally (gray matter thickness and myelination) distinct from the two sulci surrounding it: the cSTS3 and lTOS (**Fig. 1**). An rm-ANOVA (within-participant factors: sulcus, metric, and hemisphere for standardized metric units) revealed a sulcus x metric interaction (F(4, 276.19) = 179.15, η2 = 0.38, *p* < .001). Post hoc tests showed four main differences: (i) the slocs-v was shallower than cSTS3 (*p* < .001) but not lTOS (*p* = .60), (ii) the slocs-v was smaller than both the cSTS3 and lTOS (*p*s < .001), (iii) the slocs-v showed thicker gray matter than both the cSTS3 and lTOS (*p*s < .001), and iv) the slocs-v was less myelinated than both the cSTS and lTOS (*p*s < .001; **Fig. 3a**). There was also a sulcus x metric x hemisphere interaction (F(4.20, 289.81) = 4.16, η2 = 0.01, *p* = .002; hemispheric effects discussed in Supplementary Results). The morphological and architectural features of all LPC/LPOJ sulci are described in Supplementary Fig. 6.

**Fig. 3.**
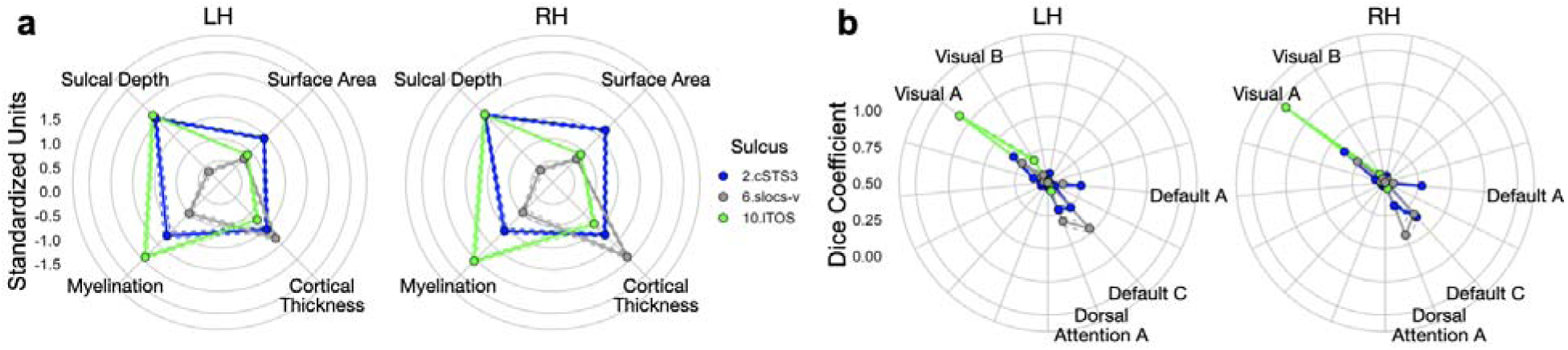
The slocs-v is morphologically, architecturally, and functionally dissociable from nearby sulci. **a.** Radial plot displaying the morphological (upper metrics: depth, surface area) and architectural (lower metrics: cortical thickness, myelination) features of the slocs-v (gray), cSTS3 (blue), and lTOS (green). Each dot and solid line represents the mean. The dashed lines indicate ± standard error. These features are colored by sulcus (legend). Metrics are in standardized units. **b.** Radial plot displaying the connectivity fingerprints of these three sulci: the Dice Coefficient overlap (values from 0-1) between each component and individual-level functional connectivity parcellations (Kong et al., 2019).

We then tested whether the slocs-v was also functionally distinct from the cSTS3 and lTOS by leveraging resting-state network parcellations for each individual participant to quantify “connectivity fingerprints” for each sulcus in each hemisphere of each participant (**Methods**; Kong et al., 2019). An rm-ANOVA (within-participant factors: sulcus, network, and hemisphere for Dice coefficient overlap) revealed a sulcus x network interaction (F(32, 2144) = 80.18, η2 = 0.55, *p* < .001). Post hoc tests showed that this interaction was driven by four effects: (i) the cSTS3 overlapped more with the Default A subnetwork than both the slocs-v and lTOS (*p*s < .001), (ii) the slocs-v overlapped more with the Default C subnetwork than the lTOS (*p* < .001) and marginally than the cSTS3 (*p* = .077), (iii) the slocs-v overlapped more with the Dorsal Attention A subnetwork than both the cSTS3 and lTOS (*p*s < .001), and iv) the lTOS overlapped more with the Visual A and Visual B subnetworks than both the cSTS3 and slocs-v (*ps* < .004; **Fig. 3b**). There was also a sulcus x network x hemisphere interaction (F(32, 2144) = 3.99, η2 = 0.06, *p* < .001; hemispheric effects discussed in Supplementary Results). Together, these results indicate that the slocs-v is a morphologically, architecturally, and functionally distinct structure from its sulcal neighbors, and thus, deserves a distinct neuroanatomical definition.

We further found that the three caudal STS rami (Petrides, 2019; Segal and Petrides, 2012) and intermediate parietal sulci (aipsJ and pips; Petrides, 2019; Zlatkina and Petrides, 2014) are morphologically, architecturally, and functionally distinct structures for the first time (to our knowledge), which empirically supports their distinctions with separate sulcal labels (Supplementary Results and Supplementary Fig. 7).

### The morphology of LPC/LPOJ sulci, including the slocs-v, is related to cognitive performance

Finally, leveraging a data-driven approach of cross-validated LASSO feature selection, we sought to determine whether sulcal depth, a main defining feature of sulci, related to cognitive performance (**Methods**). To do so, we primarily focused on spatial orientation and reasoning given that these abilities recruit multiple subregions of lateral parietal and/or occipital cortices (Gur et al., 2000; Karnath, 1997; Vendetti and Bunge, 2014; Wendelken, 2014). As in prior work (Maboudian et al., 2024; Voorhies et al., 2021; Willbrand et al., 2023b; Yao et al., 2022), we chose the model at the alpha that minimized MSE_cv_. Participants with a slocs-v in both hemispheres and all behavioral metrics were included (N = 69). Due to their rarity (being in less than 70% of hemispheres at most), we did not include the slocs-d or pAng components in this analysis.

This method revealed an association between spatial orientation scores and normalized sulcal depth in the left hemisphere (MSE_cv_ = 25.63, alpha = 0.05; **Fig. 4a**), but not in the right hemisphere (MSE_cv_ = 26.41, alpha = 0.3). Further, we found that no LPC/LPOJ sulci were selected for reasoning in either hemisphere (right: alpha = 0.3, MSE = 24.01; left: alpha = 0.3, MSE = 24.01). Six left hemisphere LPC/LPOJ sulci were related to spatial orientation task performance (**Fig. 4a, b**). Four of these sulci were positioned ventrally: cSTS3 (β = −9.77), slocs-v (β = −3.36), lTOS (β = −4.91), and mTOS (β = −0.06), whereas two were positioned dorsally: pips (β = 5.02), and SPS (β = 4.30; **Fig. 4a, b**). Using LooCV to construct models that predict behavior, the LASSO-selected model explained variation in spatial orientation score (R^2^_cv_ = 0.06, MSE_cv_ = 23.99) above and beyond a model with all left hemisphere sulci (R^2^ < 0.01, MSE_cv_ = 27.12). This model also showed a moderate correspondence (r_s_ = 0.29, p = .01; **Fig. 4c**) between predicted and actual measured scores. We then tested for anatomical and behavioral specificity using the AIC, which revealed two primary findings. First, we found that the LASSO-selected sulcal depth model outperformed a model using the cortical thickness of the six LASSO-selected sulci (R^2^_cv_ < .01, MSE_cv_ = 26.02, AIC_cortical thickness_ – AIC_sulcal depth_ = 2.19). This model also showed task specificity as these sulci outperformed a model with processing speed (R^2^_cv_ < .01, MSE_cv_ = 254.65, AIC_processing speed_ – AIC_spatial orientation_ = 63.57). Thus, our data-driven model explains a significant amount of variance on a spatial orientation task and shows behavioral and morphological specificity.

**Fig. 4.**
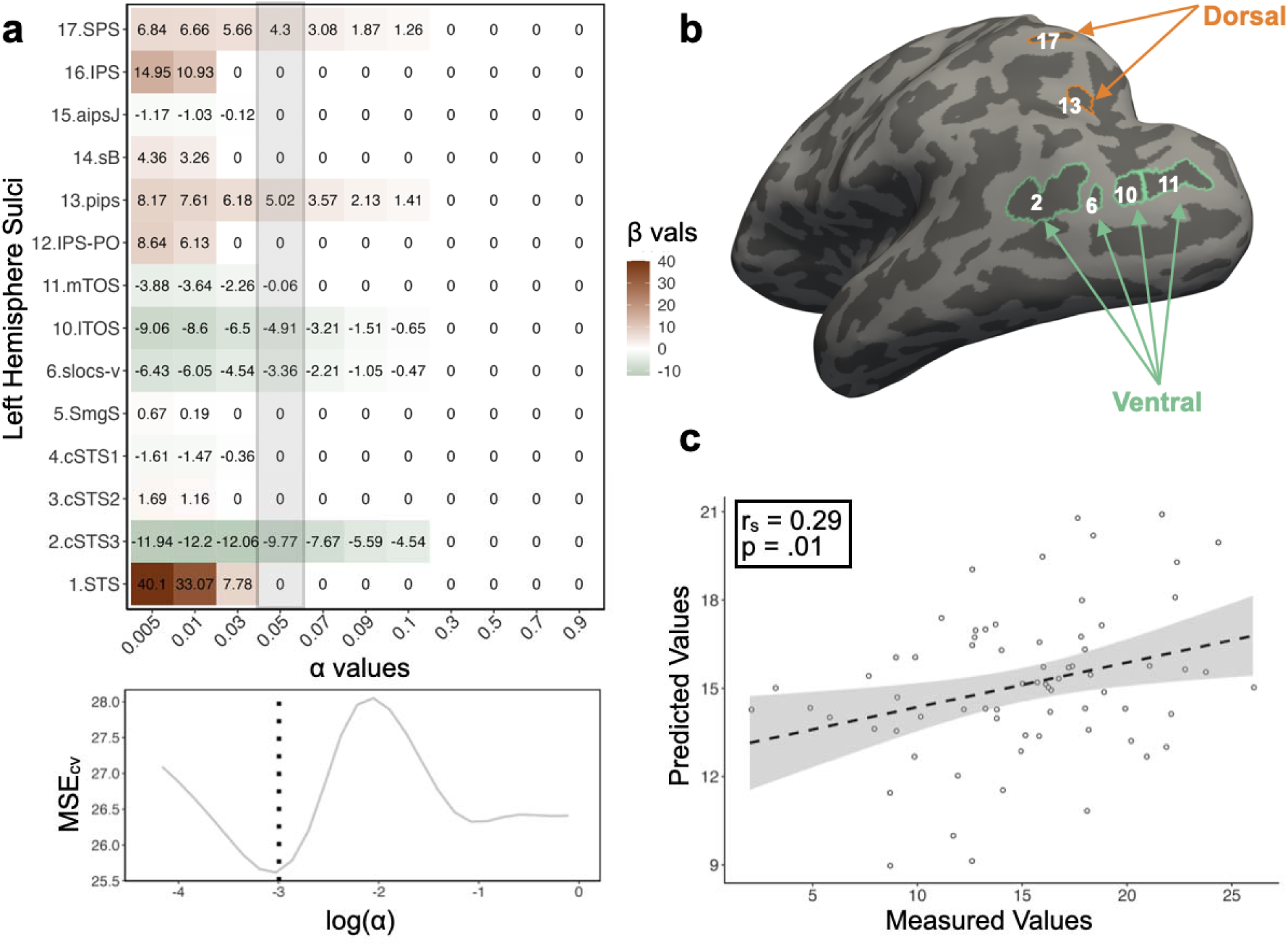
The morphology of LPC/LPOJ sulci, including the slocs-v, is related to cognitive performance. **a.** Beta-coefficients for each left hemisphere LPC/LPOJ sulcus at a range of shrinking parameter values (alpha, α). The highlighted gray bar indicates coefficients at the chosen α-level. Bottom: Cross-validated mean-squared error (MSE_CV_) at each α level. By convention, we selected the α that minimized the MSE_CV_ (dotted line). **b.** Inflated left hemisphere cortical surface from an example participant highlighting the two groups of sulci—*dorsal positive* (orange) and *ventral negative* (green)—related to spatial orientation performance. **c.** Spearman’s correlation (r_s_) between the measured and the predicted spatial orientation scores from the LASSO-selected model is shown in a.

Finally, as in prior work examining variably-present PTS in other cortical expanses (for example, Amiez et al., 2018; Cachia et al., 2014; Fornito et al., 2004; Willbrand et al., 2024b), we assessed whether the presence/absence of the more variable PTS identified in the present work (slocs-d, pAngs-v, and pAngs-d) was related to spatial orientation, reasoning, and processing speed task performance. We identified no significant associations between the presence/absence of these sulci in either hemisphere with performance on these tests (*p*s > .05).

## Discussion

### Overview

In the present study, we examined the relationship between LPC/LPOJ sulcal morphology, functional connectivity fingerprints, and cognition. We report five main findings. First, while manually defining sulci in LPC/LPOJ across 144 hemispheres, we uncovered four small and shallow sulci that are not included in present or classic neuroanatomy atlases or neuroimaging software packages. Second, we found that the most common of these structures (the slocs-v; identifiable 98.6% of the time) was morphologically, architecturally, and functionally differentiable from nearby sulci. Third, using a model-based, data-driven approach quantifying the relationship between sulcal morphology and cognition, we found a relationship between the depths of six LPC/LPOJ sulci and performance on a spatial orientation processing task. Fourth, the model identified distinct dorsal and ventral sulcal networks in LPC/LPOJ: ventral sulci had negative weights while dorsal sulci had positive weights (**Fig. 4b**). These findings are consistent with previous neuroimaging work from Gur et al. (2000) who demonstrated separate functional activations in dorsal parietal and the more ventrally situated occipital-parietal cortices for the judgment of line orientation task used in the present study. Fifth, the model identified that the slocs-v is cognitively relevant, further indicating the importance of this neuroanatomical structure. In the sections below, we discuss (i) the slocs-v relative to modern functional and cytoarchitectonic parcellations in the LPC/LPOJ, as well as anatomical connectivity to other parts of the brain, (ii) underlying anatomical mechanisms relating sulcal morphology and behavior more broadly, and (iii) limitations of the present study. Implications for future studies are distributed throughout each section.

### The slocs-v relative to modern functional and cytoarchitectonic parcellations in the LPC/LPOJ, as well as anatomical connectivity to other parts of the brain

To lay the foundation for future studies relating the newly-described slocs-v to different anatomical and functional organizational features of LPC/LPOJ, we situate probabilistic predictions of slocs-v relative to probabilistic cortical areas identified using multiple modalities. For example, when examining the correspondence between the slocs-v and modern multimodal (Human Connectome Project multimodal parcellation, HCP-MMP; Glasser et al., 2016) and observer-independent cytoarchitectural (Julich-Brain atlas; Amunts et al., 2020) areas (**Methods**), the slocs-v is located within distinct areas. In particular, the slocs-v aligns with the multimodally- and cytoarchitecturally-defined area PGp bilaterally and cytoarchitecturally-defined hIP4 in the right hemisphere (**Fig. 5**). In classic neuroanatomical terms (Cunningham, 1892), this indicates that the slocs-v is a putative “axial sulcus” for these regions, which future work can assess with analyses in individual participants.

**Fig. 5.**
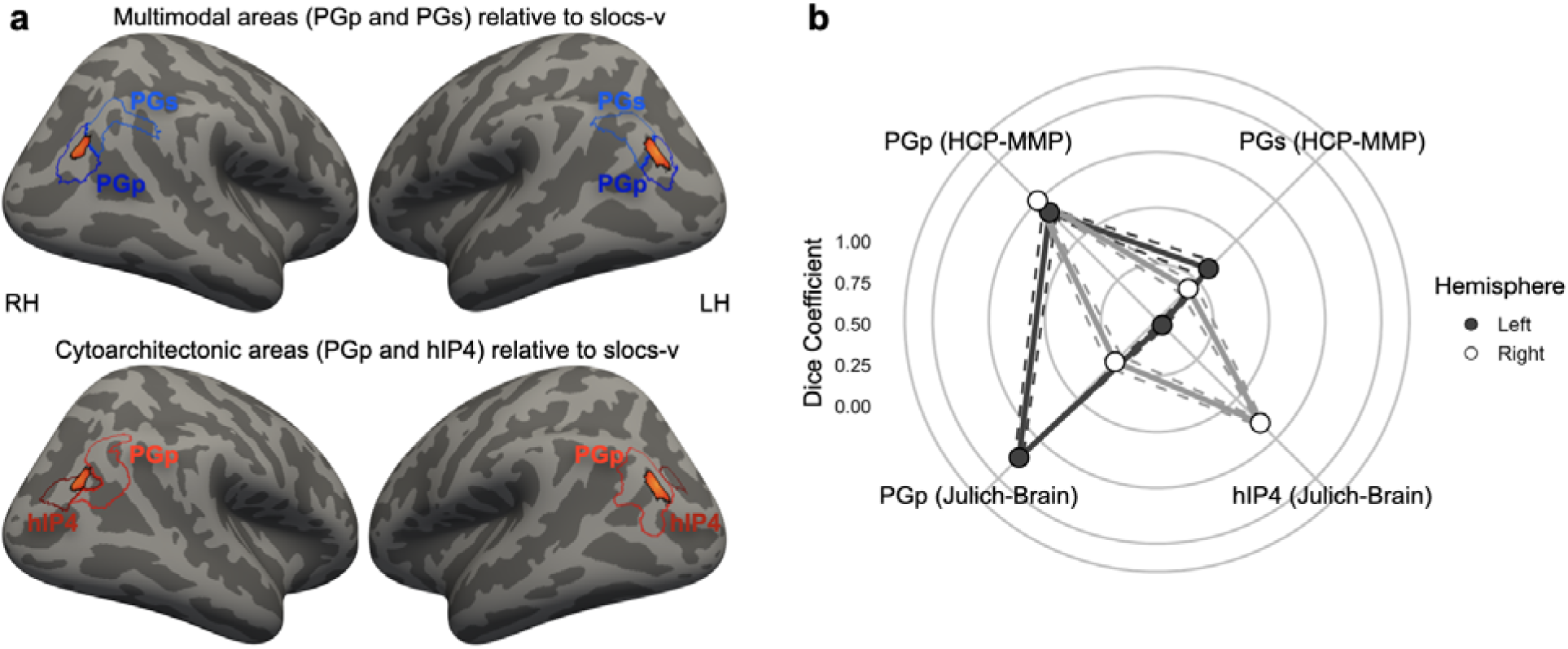
The slocs-v relative to modern functional and cytoarchitectonic parcellations in LPC/LPOJ. **a.** Top: Left (LH) and right (RH) hemispheres of the inflated fsaverage surface with two areas from the modern HCP multimodal parcellation (HCP-MMP; blue; Glasser et al., 2016) relative to an MPM of the slocs-v (warm colors indicate areas with at least 20% overlap across participants; Fig. 2). Bottom: Same as top, except for two observer-independent cytoarchitectonic regions from the Julich-Brain Atlas (Amunt et al., 2020). **b.** Overlap between the slocs-v and each area (Methods). Each dot and solid line represents the mean. The dashed lines indicate ± standard error (left: gray; right: white).

Aside from recent multimodal and observer-independent cytoarchitectonic parcellations, an immediate question is: What is the relationship between the slocs-v and other functional regions at this junction between the occipital and parietal lobes, as well as potential anatomical connectivity? For example, there are over a dozen visual field maps in the cortical expanse spanning the TOS, IPS-PO, and the IPS proper (see (i), (ii), and (iii), respectively in **Fig. 6a**; Mackey et al., 2017). When projecting probabilistic locations of retinotopic maps from over 50 individuals from Wang and colleagues (Wang et al., 2015; **Methods**), the slocs-v is likely located outside of visual field maps extending into this cortical expanse (**Fig. 6a**). Nevertheless, when also projecting the map of the mean R^2^ metric from the HCP retinotopy dataset from 181 participants shared by Benson and colleagues (Benson et al., 2018; **Methods**), the slocs-v is in a cortical expanse that explains a significant amount of variance (left hemisphere: R^2^ = 19.29, R^2^_max_ = 41.73; right hemisphere: R^2^_mean_ = 21.17, R^2^_max_ = 44.23; **Fig. 6b**).

**Fig. 6.**
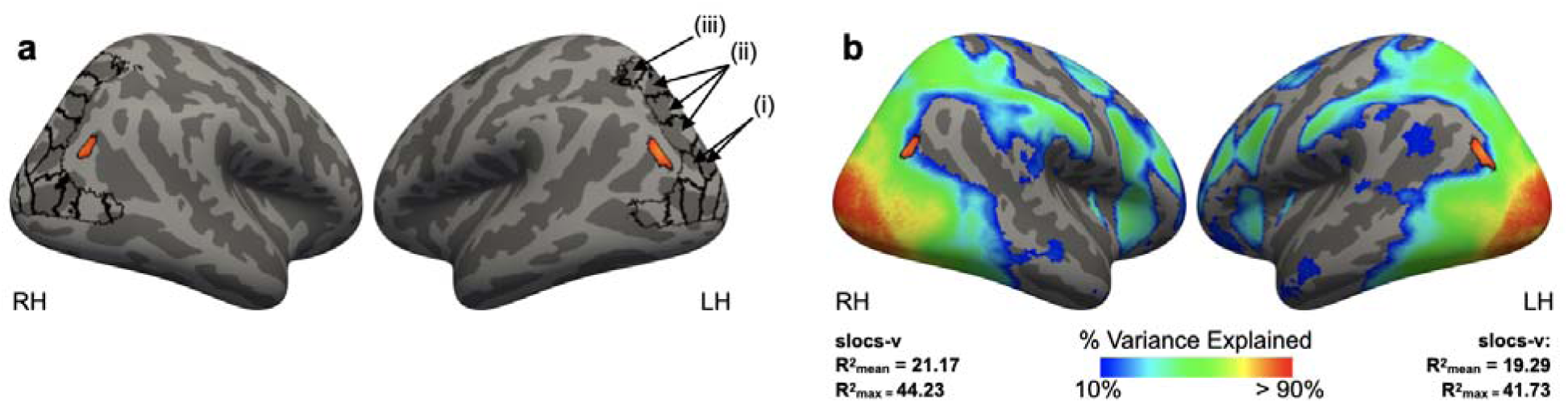
The slocs-v relative to retinotopy. **a.** Top: Left (LH) and right (RH) hemispheres of the inflated fsaverage surface displaying the probabilistic locations of retinotopic maps from over 50 individuals from Wang and colleagues (Wang et al., 2015); black outlines). The predicted slocs-v location from the MPM is overlaid in orange (as in Fig. 4). (i), (ii), and (iii) point out the retinotopic maps in the cortical expanse spanning the TOS, IPS-PO, and IPS, respectively. **b.** Same format as in a, but with a map of the mean R^2^ metric from the HCP retinotopy dataset (Benson et al., 2018) overlaid on the fsaverage surfaces (thresholded between R^2^ values of 10% and 90%). This metric measures how well the fMRI time series at each vertex is explained by a population receptive field (pRF) model. The mean and max R^2^ values for the slocs-v MPM in each hemisphere are included below each surface.

In terms of anatomical connectivity, as the slocs-v co-localizes with cytoarchitectonically defined PGp (**Fig. 5**) and previous studies have examined the anatomical connectivity of the probabilistically defined PGp, we can glean insight regarding the anatomical connectivity of slocs-v from these previous studies (Caspers et al., 2011; Wang et al., 2012). This prior work showed that PGp was anatomically connected to temporo-occipital regions, other regions in the temporal lobe, middle and superior frontal cortex, as well as the inferior frontal cortex and insula (Caspers et al., 2011; Wang et al., 2012). Furthermore, the slocs-v appears to lie at the junction of scene-perception and place-memory activity (a transition that also consistently co-localizes with the HCP-MMP area PGp) as identified by Steel and colleagues (2021). Of course, the location of the slocs-v relative to multimodal, cytoarchitectonic, and retinotopic areas, as well as the anatomical connectivity of the slocs-v, would need to be examined in individual participants, but the present work makes clear predictions for future studies as fleshed out here. To conclude this section, as the multimodal area PGp (**Fig. 5**) was recently proposed as a “transitional area” by Glasser and colleagues (2016; Supplementary Table 5), future studies can also further functionally and anatomically test the transitional properties of slocs-v.

### Underlying anatomical mechanisms relating sulcal morphology and behavior

In this section, we discuss potential anatomical mechanisms contributing to the relationship between sulcal depth and behavior in two main ways. First, long-range white matter fibers have a gyral bias, while short-range white matter fibers have a sulcal bias in which some fibers project directly from the deepest points of a sulcus (Cottaar et al., 2021; Reveley et al., 2015; Schilling et al., 2018, 2023; Van Essen et al., 2014). As such, recent work hypothesized a close link between sulcal depth and short-range white matter properties (Bodin et al., 2021; Pron et al., 2021; Voorhies et al., 2021; Willbrand et al., 2023b; Yao et al., 2022): deeper sulci would reflect even shorter short-range white matter fibers, which would result in faster communication between local, cortical regions and in turn, contribute to improved cognitive performance. This increased neural efficiency could underlie individual differences in cognitive performance. Ongoing work is testing this hypothesis which can be further explored in future studies incorporating anatomical, functional, and behavioral measures, as well as computational modeling.

Second, our model-based approach identified separate dorsal and ventral sulcal networks in which deeper sulci dorsally and shallower sulci ventrally contributed to the most explained variance on the spatial orientation task. A similar finding was identified by our previous work in the lateral prefrontal cortex (Yao et al., 2022). These previous and present findings may be explained by the classic anatomical compensation theory, which proposes that the size and depth of a sulcus counterbalance those of the neighboring sulci (Armstrong et al., 1995; Connolly, 1950; Zilles et al., 2013). Thus, a larger, deeper sulcus would be surrounded by sulci that are smaller and shallower, rendering the overall degree of cortical folding within a given region approximately equal (Armstrong et al., 1995; Connolly, 1950; Zilles et al., 2013). Future work can incorporate underlying white matter architecture into the compensation theory, as well as a recent modification that proposed to also incorporate local morphological features such as the deepest sulcal point (for example, sulcal pit or sulcal root; Régis et al., 2005), which has recently been shown to be related to different functional features of the cerebral cortex (Bodin et al., 2018; Leroy et al., 2015; Natu et al., 2021). Altogether, these and recent findings begin to build a multimodal mechanistic neuroanatomical understanding underlying the complex relationship between sulcal depth and cognition relative to other anatomical features.

### Limitations

The main limitation of our study is that presently, the most accurate methodology to define sulci —especially the small, shallow, and variable PTS—requires researchers to manually trace each structure on the cortical surface reconstructions. This method is limited due to the individual variability of cortical sulcal patterning (**Fig. 1**, Supplementary Fig. 5), which makes it challenging to identify sulci without extensive experience and practice. However, we anticipate that our probabilistic maps (**Fig. 2**) will provide a starting point and hopefully, expedite the identification of these sulci in new participants. This should accelerate the process of subsequent studies confirming the accuracy of our updated schematic of LPC/LOPJ. This manual method is also arduous and time-consuming, which, on the one hand, limits the sample size in terms of number of participants, while on the other, results in thousands of precisely defined sulci. This push-pull relationship reflects a broader conversation in the human brain mapping and cognitive neuroscience fields between a balance of large N studies and “precision imaging” studies in individual participants (Gratton et al., 2022; Naselaris et al., 2021; Rosenberg and Finn, 2022). Though our sample size is comparable to other studies that produced reliable results relating sulcal morphology to brain function and cognition (for example, Cachia et al., 2021; Garrison et al., 2015; Lopez-Persem et al., 2019; Miller et al., 2021; Roell et al., 2021; Voorhies et al., 2021; Weiner, 2019; Willbrand et al., 2022a, 2022b; Yao et al., 2022), ongoing work that uses deep learning algorithms to automatically define sulci should result in much larger sample sizes in future studies (Borne et al., 2020; Lee et al., 2024, 2025; Lyu et al., 2021). The time-consuming manual definitions of primary, secondary, and PTS also limit the cortical expanse explored in each study, thus restricting the present study to LPC/LPOJ.

Additionally, the scope of the present study is limited in that the sample was only in young adults. This sample was selected as it is the standard of the field when charting features of PTS for the first time (for example, Chiavaras and Petrides, 2000; Drudik et al., 2023; Miller et al., 2021; Paus et al., 1996; Segal and Petrides, 2012; Sprung-Much and Petrides, 2018; Willbrand et al., 2023c, 2022a; Zlatkina and Petrides, 2014). Nevertheless, it is necessary to explore how well this updated sulcal landscape translates to different age groups, species, and clinical populations.

It is also worth noting that the morphological-behavioral relationship identified in the present study explains a modest amount of variance; however, the more important aspect of our findings is that multiple sulci identified in our model-based approach are recently-characterized sulci in LPC/LOPJ identified by our group and others (Petrides, 2019), and thus, the relationship would have been overlooked or lost if these sulci were not identified. Finally, although we did not focus on the relationship between the other three PTS (slocs-d, pAngs-v, and pAngs-d) to anatomical and functional features of LPC and cognition, given that variability in sulcal incidence is cognitively (for review see, Cachia et al., 2021), anatomically (Amiez et al., 2021; Vogt et al., 1995), functionally (Lopez-Persem et al., 2019), and translationally (Clark et al., 2010; Le Provost et al., 2003; Meredith et al., 2012; Nakamura et al., 2020; Yücel et al., 2003, 2002) relevant, future work can also examine the relationship between the more variable slocs-d, pAngs-v, and pAngs-d and these features.

### Conclusion

In conclusion, we uncovered four previously-undefined sulci in LPC/LPOJ and quantitatively showed that the slocs-v is a stable sulcal landmark that is morphologically, architecturally, and functionally differentiable from surrounding sulci. We further used a data-driven, model-based approach relating sulcal morphology to behavior, which identified different relationships of ventral and dorsal LPC/LPOJ sulcal networks contributing to the perception of spatial orientation. The model identified the slocs-v, further indicating the importance of this newly-described neuroanatomical structure. Altogether, this work provides a scaffolding for future “precision imaging” studies interested in understanding how anatomical and functional features of LPC/LPOJ relate to cognitive performance at the individual level.

## Methods

### Participants

Data for the young adult human cohort analyzed in the present study were from the Human Connectome Project (HCP) database (https://www.humanconnectome.org/study/hcp-young-adult/overview). Here, we used 72 randomly-selected participants, balanced for gender (following the terminology of the HCP data dictionary), from the HCP database (50% female, 22-36 years old, and 90% right-handed; there was no effect of handedness on our behavioral tasks; Supplementary Methods) that were also analyzed in several prior studies (Hathaway et al., 2023; Maboudian et al., 2024; Miller et al., 2021, 2020; Willbrand et al., 2024a, 2023b, 2023c, 2022a). HCP consortium data were previously acquired using protocols approved by the Washington University Institutional Review Board (Mapping the Human Connectome: Structure, Function, and Heritability; IRB # 201204036). Informed consent was obtained from all participants.

### Neuroimaging data acquisition

Anatomical T1-weighted (T1-w) MRI scans (0.8 mm voxel resolution) were obtained in native space from the HCP database. Reconstructions of the cortical surfaces of each participant were generated using FreeSurfer (v6.0.0), a software package used for processing and analyzing human brain MRI images (<u>surfer.nmr.mgh.harvard.edu</u>; Dale et al., 1999; Fischl et al., 1999). All subsequent sulcal labeling and extraction of anatomical metrics were calculated from these native space reconstructions generated through the HCP’s version of the FreeSurfer pipeline (Glasser et al., 2013).

### Behavioral data

In addition to structural and functional neuroimaging data, the HCP also includes a wide range of behavioral metrics from the NIH toolbox (Barch et al., 2013). To relate LPC/LPOJ sulcal morphology to behavior, we leveraged behavioral data related to spatial orientation (Variable Short Penn Line Orientation Test), relational reasoning (Penn Progressive Matrices Test), and processing speed (Pattern Completion Processing Speed Test; Supplementary Methods for task details). We selected these tasks as previous functional neuroimaging studies have shown the crucial role of LPC/LPOJ in relational reasoning and spatial orientation (Gur et al., 2000; Karnath, 1997; Vendetti and Bunge, 2014; Wendelken, 2014), while our previous work relating sulcal morphology to cognition uses processing speed performance as a control behavioral task (Voorhies et al., 2021; Willbrand et al., 2024a, 2022b).

### Anatomical analyses

#### Manual labeling of LPC sulci

Sulci were manually defined in 72 participants (144 hemispheres) guided by the most recent atlas by Petrides (2019), as well as recent empirical studies (Drudik et al., 2023; Segal and Petrides, 2012; Zlatkina and Petrides, 2014), which together offer a comprehensive definition of cerebral sulcal patterns, including PTS. For a historical analysis of sulci in this cortical expanse, please refer to Segal & Petrides (2012) and Zlatkina & Petrides (2014). Our cortical expanse of interest was bounded by the following sulci and gyri: (i) the postcentral sulcus (PoCS) served as the anterior boundary, (ii) the superior temporal sulcus (STS) served as the inferior boundary, (iii) the superior parietal lobule (SPL) served as the superior boundary, and (iv) the medial and lateral transverse occipital sulci (mTOS and lTOS) served as the posterior boundary. We also considered the following sulci within this cortical expanse: the three different branches of the caudal superior temporal sulcus (posterior to anterior: cSTS3, 2, 1), the supramarginal sulcus (SmgS), posterior intermediate parietal sulcus (pips), sulcus of Brissaud (sB), anterior intermediate parietal sulcus of Jensen (aipsJ), paroccipital intraparietal sulcus (IPS-PO), intraparietal sulcus (IPS), and the superior parietal sulcus (SPS). Of note, the IPS-PO is the portion of the IPS extending ventrally into the occipital lobe. The IPS-PO was first identified as the paroccipital sulcus by Wilder (1886). There is often an annectant gyrus separating the horizontal portion of the IPS proper from the IPS-PO (Roell et al., 2021; Zlatkina and Petrides, 2014).

Additionally, we identified as many as four previously uncharted and variable LPC/LPOJ PTS for the first time: the supralateral occipital sulcus (slocs; composed of ventral (slocs-v) and dorsal (slocs-d) components) and the posterior angular sulcus (pAngs; composed of ventral (pAngs-v) and dorsal (pAngs-d) components). In the Supplementary Methods and Supplementary Figs. 1–4, we discuss the slocs and pAngs within the context of modern and historical sources.

As this is the first time the sulcal expanse of LPC/LOPJ was comprehensively charted with a focus on pTS, the location of each sulcus was confirmed through a three-tiered procedure for each participant in each hemisphere. First, trained independent raters (Y.T. and T.G.) identified sulci. Second, these definitions were checked by a trained expert (E.H.W.). Third, these labels were finalized by a neuroanatomist (K.S.W.). We emphasize that this procedure has produced reproducible results in our prior work across the cortex (Hastings et al., 2024; Maboudian et al., 2024; Miller et al., 2021, 2020; Parker et al., 2023; Ramos Benitez et al., 2024; Voorhies et al., 2021; Willbrand et al., 2024a, 2024b, 2023b, 2023c, 2022a, 2022b; Yao et al., 2022). All LPC sulci were then manually defined and saved as .label files in FreeSurfer using tksurfer tools, from which morphological and anatomical features were extracted. We defined LPC/LPOJ sulci for each participant based on the most recent schematics of sulcal patterning by Petrides (2019) as well as pial, inflated, and smoothed white matter (smoothwm) FreeSurfer cortical surface reconstructions of each individual. In some cases, the precise start or end point of a sulcus can be difficult to determine on a surface (Borne et al., 2020); however, examining consensus across multiple surfaces allowed us to clearly determine each sulcal boundary in each individual. For four example hemispheres with these 13-17 sulci identified, see **Fig. 1a** (Supplementary Fig. 5 for all hemispheres). The specific criteria to identify the slocs and pAngs are outlined in **Fig. 1b**.

To test whether the incidence rates of the slocs and pAngs components were statistically different, we implemented a binomial logistic regression GLM with sulcus (slocs-v, slocs-d, pAngs-v, and pAngs-d) and hemisphere (left and right), as well as their interaction, as predictors for sulcal presence (0: absent, 1: present). Additional GLMs were run relating the incidence of the more variable sulci (slocs-d, pAngs-v, and pAngs-d) to demographic features (gender and age) were also run. GLMs were carried out with the glm function from the built-in stats R package. ANOVA χ2 tests were applied to each GLM with the Anova function from the car R package, from which results were reported.

### Probability maps

Sulcal probability maps were generated to show the vertices with the highest alignment across participants for a given sulcus. To create these maps, the label file for each sulcus was transformed from the individual to the fsaverage surface with the FreeSurfer mri_label2label command (https://surfer.nmr.mgh.harvard.edu/fswiki/mri_label2label). Once each label was transformed into this common template space, we calculated the proportion of participants for which each vertex was labeled as the given sulcus with custom Python code (Miller et al., 2021; Voorhies et al., 2021). For vertices with overlap between sulci, we employed a “winner-take-all” approach such that the sulcus with the highest overlap across participants was assigned to that vertex. Alongside the thresholded maps, we also provide constrained maps (maximum probability maps, MPMs) at 20% participant overlap to increase interpretability (20% MPMs shown in **Fig. 2**). To aid future studies interested in investigating LPC/LPOJ sulci, we share these maps with the field (**Data availability**).

### Extracting and comparing the morphological and architectural features from sulcal labels

Morphologically, we compared sulcal depth and surface area across sulci, as these are two of the primary morphological features used to define and characterize sulci (Armstrong et al., 1995; Chi et al., 1977; Leroy et al., 2015; Lopez-Persem et al., 2019; Miller et al., 2021, 2020; Natu et al., 2021; Sanides, 1964; Voorhies et al., 2021; Weiner, 2019; Welker, 1990; Willbrand et al., 2023b, 2022a; Yao et al., 2022). As in our prior work (Voorhies et al., 2021; Yao et al., 2022), mean sulcal depth values (in standard FreeSurfer units) were computed in native space from the .sulc file generated in FreeSurfer (Dale et al., 1999) with custom Python code (Voorhies et al., 2021). Briefly, depth values are calculated based on how far removed a vertex is from what is referred to as a “mid-surface,” which is determined computationally so that the mean of the displacements around this “mid-surface” is zero. Thus, generally, gyri have negative values, while sulci have positive values. Each depth value was also normalized by the deepest point in the given hemisphere. Surface area (mm^2^) was calculated with the FreeSurfer mris_anatomical_stats function (https://surfer.nmr.mgh.harvard.edu/fswiki/mris_anatomical_stats). The morphological features of all LPC/LPOJ sulci are documented in Supplementary Fig. 6.

Architecturally, we compared cortical thickness and myelination, as in our prior work in other cortical expanses (Miller et al., 2021; Voorhies et al., 2021; Willbrand et al., 2023b, 2022a). Mean gray matter cortical thickness (mm) was extracted using the FreeSurfer mris_anatomical_stats function. To quantify myelin content, we used the T1-w/T2-w maps for each hemisphere, an in vivo myelination proxy (Glasser and Van Essen, 2011). To generate the T1-w/T2-w maps, two T1-w and T2-w structural MR scans from each participant were registered together and averaged as part of the HCP processing pipeline (Glasser et al., 2013). The averaging helps to reduce motion-related effects or blurring. Additionally, and as described by Glasser and colleagues (2013), the T1-w/T2-w images were bias-corrected for distortion effects using field maps. We then extracted the average T1-w/T2-w ratio values across each vertex for each sulcus using custom Python code (Miller et al., 2021). The architectural features of all LPC/LPOJ sulci are documented in Supplementary Fig. 6.

To assess whether these four metrics differed between the slocs-v and surrounding sulci (cSTS3 and lTOS), we ran a repeated measure analysis of variance (rm-ANOVA) with the within-participant effects of sulcus (slocs-v, cSTS3, and lTOS), metric (surface area, depth, cortical thickness, and myelination), and hemisphere (left and right). Rm-ANOVAs (including sphericity correction) were implemented with the aov_ez function from the afex R package. Effect sizes for the ANOVAs are reported with the partial eta-squared metric (η2). Post-hoc analyses were computed with the emmeans function from the emmeans R package (*p*-values corrected with Tukey’s method). We also repeated these analyses for the three cSTS components (Petrides, 2019; Segal and Petrides, 2012) and the two intermediate parietal sulcal components (ips: aipsJ and pips; Petrides, 2019; Zlatkina and Petrides, 2014; detailed in the Supplementary Results and Supplementary Fig. 7) as these components, to our knowledge, have not been quantitatively compared in previous work.

### Functional analyses

To determine if the slocs-v is functionally distinct from surrounding sulci, we generated functional connectivity profiles using recently developed analyses (Miller et al., 2021; Willbrand et al., 2023a, 2022a). First, we used resting-state network parcellations for each individual participant from Kong and colleagues (2019), who generated individual network definitions by applying a hierarchical Bayesian network algorithm to produce maps for each of the 17 networks in individual HCP participants. Importantly, this parcellation was conducted blind to both cortical folding and our sulcal definitions. Next, we resampled the network profiles for each participant onto the fsaverage cortical surface, and then to each native surface using CBIG tools (https://github.com/ThomasYeoLab/CBIG). We then calculated the spatial overlap between a sulcus and each of the 17 individual resting-state networks via the Dice coefficient (Equation 1):

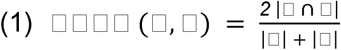

This process of calculating the overlap between each sulcus and the 17-network parcellation generated a “connectivity fingerprint” for each sulcus in each hemisphere of each participant. We then ran an rm-ANOVA with within-participant factors of sulcus (slocs-v, cSTS3, and lTOS), network (17 networks), and hemisphere (left and right) to determine if the network profiles (i.e., the Dice coefficient overlap with each network) of the slocs-v was differentiable from the surrounding sulci (i.e., cSTS3 and lTOS). Here we discuss effects related to networks that at least showed minor overlap with one sulcus (i.e., Dice ≥ .10). As in the prior analysis, we also repeated these analyses for the three cSTS components and the two intermediate parietal sulcal components (Supplementary Results and Supplementary Fig. 7).

### Behavioral analyses

#### Model selection

The analysis relating sulcal morphology to spatial orientation and/or reasoning consisted of using a cross-validated (CV) least absolute shrinkage and selection operator (LASSO) regression to select the sulci that explained the most variance in the data and determined how much variance is explained by sulcal depth as a predictor of behavior, as implemented in our previous work (Maboudian et al., 2024; Voorhies et al., 2021; Willbrand et al., 2023b; Yao et al., 2022). A LASSO regression is well suited to address our question since it facilitates the model selection process and increases the generalizability of a model by providing a sparse solution that reduces coefficient values and decreases variance in the model without increasing bias (Heinze et al., 2018). Further, regularization is recommended in cases where there are many predictors (X > 10), as in this study, because this technique guards against overfitting and increases the likelihood that a model will generalize to other datasets. A LASSO performs L1 regularization by applying a penalty, or shrinking parameter (alpha, α), to the absolute magnitude of the coefficients. In this manner, low coefficients are set to zero and eliminated from the model. Therefore, LASSO affords data-driven variable selection that results in simplified models containing only the most predictive features, in this case, sulci predicting cognitive performance. This methodology improves model interpretability and prediction accuracy, as well as protects against overfitting, which improves generalizability (Ghojogh and Crowley, 2019; Heinze et al., 2018).

The depths of all LPC/LPOJ sulci were included as predictors in the LASSO regression model (Supplementary Methods for details on demographic control variables). We used nested CV to optimize the shrinking parameter for the LASSO regression. By convention (Heinze et al., 2018), we selected the model parameters that minimized the CV mean squared error (MSE_cv_). Optimization was performed with the GridSearchCV function from the SciKit-learn package in Python. This function allowed us to determine the model parameters minimizing the MSE_cv_ by performing an exhaustive search across a range of α values. Nested CV was done as non-nested CV leads to biased performance (Cawley and Talbot, 2010; Vabalas et al., 2019).

To evaluate the performance of the model selected by the LASSO regression and verify the result of our feature selection, we used linear regression with leave-one-out CV (LooCV) to fit these selected models and to compare various models. Specifically, we measured the model performance for the relevant behavioral task using nested model comparison. With LooCV, we compared the LASSO-selected model with the predictors to a model with all left hemisphere sulci as predictors. All regression models were implemented with functions from the SciKit-learn Python package.

### Assessing morphological and behavioral specificity

To assess whether our findings generalized to other anatomical features, we considered cortical thickness, which is consistently studied in cognitive neuroscience studies relating morphology to cognition (Dickerson et al., 2008; Gogtay et al., 2004; Maboudian et al., 2024; Voorhies et al., 2021; Willbrand et al., 2023b; Yao et al., 2022). To do so, we replaced sulcal depth with cortical thickness as the predictive metric in our LASSO-selected model. As with depth, the model was fit to the data with LooCV. To compare the thickness model to the depth model, we used the Akaike Information Criterion (AIC), which provides an estimate of in-sample prediction error and is suitable for non-nested model comparison. By comparing AIC scores, we are able to assess the relative performance of the two models. If the ΔAIC is > 2, it suggests an interpretable difference between models. If the ΔAIC is > 10, it suggests a strong difference between models, with the lower AIC value indicating the preferred model (Wagenmakers and Farrell, 2004). To also ascertain whether the relationship between LPC/LPOJ sulcal depth and cognition is specific to spatial orientation performance, or transferable to other general measures of cognitive processing, we investigated the generalizability of the sulcal-behavior relationship to another widely used measure of cognitive functioning: processing speed (Kail and Salthouse, 1994). Specifically, we used LooCV to predict processing speed instead of spatial orientation score. As with thickness, we compared the two models with the AIC.

### Assessing the relationship between variable presence of the slocs-d, pAngs-v, and pAngs-d and behavior

To test the relationship between the more variable PTS identified in the present work and behavior, we implemented t-tests to assess the presence of the slocs-d, pAngs-v, and pAngs-d in both the left and right hemisphere to spatial orientation, reasoning, and processing speed task performance.

### Situating the slocs-v within modern group-level cortical parcellations

To putatively relate the slocs-v to modern multimodal (HCP multimodal parcellation, HCP-MMP; Glasser et al., 2016) and cytoarchitectural (Julich-Brain atlas; Amunts et al., 2020) regions of the cerebral cortex located in fsaverage template space, we quantified the Dice coefficient overlap between the slocs-v of each participant (resampled to fsaverage space) and the individual regions of interest comprising the HCP-MMP and Julich-Brain parcellations.

### Retinotopic response mapping of LPC/LPOJ sulci

To assess whether any of the LPC/LPOJ sulci related to retinotopic representations, we leveraged population receptive field mapping data (Benson et al., 2018). For each sulcal MPM (as the retinotopic data were only available in this template space), we extracted the mean R^2^ values (i.e., the percentage of variance in each vertex explained by the population receptive field model) for vertices that showed meaningful retinotopic responses across participants (thresholded at R^2^ > 10%; Mackey et al., 2017).

## Supplementary Methods

### In-depth description of behavioral tasks

#### *V*ariable Short Penn Line Orientation Test

In this study, we used behavioral data related to spatial orientation from the National Institutes of Health (NIH) toolbox (Barch et al., 2013). In this toolbox, spatial orientation processing was measured as performance on the Variable Short Penn Line Orientation Test (also called the Judgment of Line Orientation Test, JOLO; Benton et al., 1975; Gur et al., 2010, 2001a, 2001b, 1982). The JOLO has been designed to evaluate the ability to match the orientation and the angle of lines in space. At first, two lines of different colors and orientations are presented to the participant. One of them (blue) must be manually rotated so that it becomes parallel with the second line (red) which remains fixed. To match the angled lines, participants have the possibility to rotate the first line either clockwise or counterclockwise. As the various trials are carried out, the lines vary in their location and distance on the screen. The line to be turned by the participant can also vary in size (long or short) while the second line remains fixed. Spatial Orientation processing is measured like so with 24 different trials.

#### Penn Progressive Matrices Test

From the same NIH toolbox (Barch et al., 2013), we also used behavioral data related to fluid intelligence (i.e., relational reasoning; Christoff et al., 2001; Conway et al., 2005; Gray et al., 2005, 2003; Prabhakaran et al., 1997; Wendelken et al., 2008). Specifically, for each Human Connectome Project (HCP) participant, relational (matrix) reasoning scores were measured as the total Penn Progressive Matrices task score from form A of the abbreviated version of the Raven’s Progressive Matrices (Bilker et al., 2012). In this task, each participant is instructed to determine the missing element that completes the matrix by selecting the right pattern among an array of options. They must make a choice in such a way that the two bottom shapes mirror the relationship between the two uppermost shapes. Participants must pick one pattern among five different options. The entire task is composed of 24 different matrices to complete, in order of increasing difficulty. However, after five incorrect choices in a row, the task discontinues.

#### Pattern Completion Processing Speed Test

From the same NIH toolbox (Barch et al., 2013), we also used behavioral data related to processing speed, tested via the Pattern Completion Processing Speed Test (Carlozzi et al., 2015). As in prior work (Voorhies et al., 2021; Willbrand et al., 2022b), this was used as a behavioral control metric. This test has been designed to measure the speed of processing based on the ability of the participant to discern whether or not two pictures that are side-by-side are the same as fast as possible. During the test, participants have to discriminate among different types of differences (addition/removal of an element or again the color or the number of elements on the pictures). The final score corresponds to the number of correct answers during a 90-seconds period. Participants’ responses are made by pressing a “yes” or “no” button.

### LPOJ sulci relative to historical atlases and modern investigations

In a series of papers, Petrides and colleagues (Segal and Petrides, 2012; Zlatkina and Petrides, 2014) discuss historical contentions regarding the caudal rami of the STS (cSTS), as well as the multiple portions of the IPS, including the aipsJ and the pips. Here, we complement their historical analyses by also incorporating additional classic sources that either depicted or attempted to label small sulci between the branches of the STS and the IPS components in the vicinity of the slocs and pAngs components identified in the present study (**Supplementary Fig. 1**), which are discussed in separate subsections below.

#### slocs vs. prelunate

The cortical expanse of interest in the present study is bounded by the caudal branches of the STS anteriorly and the body of the occipital portion of the IPS, or IPS-PO, which historically was originally labeled as the paroccipital sulcus by Wilder (1886). The slocs components should not be confused with what has been referred to as the superior occipital sulcus (SOS), which is another name for the IPS-O, or paroccipital (Kujovic et al., 2013; Malikovic et al., 2012). Three caudal branches of the STS have been identified throughout history, though as summarized by Segal and Petrides (2012), modern atlases that are extensively cited (for example, Ono et al., 1990; Duvernoy, 1999) identify two branches of the STS—and confusingly, define them differently. Nevertheless, classic anatomists consistently identified three caudal STS rami in different species: Kükenthal and Ziehen (1895) in different non-human hominoids such as orangutans and chimpanzees, Bolk (1909) in gorillas, and Connolly (1950) in each of those species, as well as in humans (**Supplementary Figs. 1, 4**).

In the latter study, Connolly (1950) labeled a prelunate sulcus (as others before him) across many species such as gibbons, orangutans, gorillas, chimpanzees, and humans (**Supplementary Fig. 4**) in which his “pl” was much more prominent in humans compared to other species. In Connolly’s depictions, his “pl” is ventral to a depicted, but unlabeled sulcus that is defined as slocs-v in the present study. In 50 example hemispheres included from work by Connolly (1950) in Supplementary Fig. 4, this sulcus was depicted, but unlabeled, 94% of the time (47/50 hemispheres). A minority of the time (4%; 2/50 hemispheres), Connolly labeled multiple branches of either “pl” or cSTS3 (as a^3^ in his depictions), in which one of these branches is slocs-v as identified in the present study.

Consistent with the more dorsal positioning of slocs-v relative to the prelunate, Segal and Petrides (2012) detail that the prelunate has been renamed the lateral occipital sulcus, which is ventral to the slocs components we identify here. Segal and Petrides (2012) write: “The LOCS or prelunate sulcus is a horizontal sulcus that extends anteriorly from the lunate sulcus (also called the sulcus prelunatus by Elliot Smith, 1907 and by Shellshear, 1927). The LOCS is found ventral to the TOCS (see Fig. 2).” pg. 2037^1^*

Contrary to this modern definition of the prelunate sulcus, or LOCS, the slocs-v is not ventral to the TOS, but situated more dorsally between cSTS3 and the TOS. In work defining sulci in the lateral portion of the occipital lobe, Iaria and Petrides (2007) sometimes labeled our slocs-d and slocs-v as “accessory” sulci, as well as left them unlabeled (**Supplementary Fig. 2**).

#### pAngs vs. sulcus intermedius primus and secundus of Eberstaller

To our knowledge, our pAngs components are independent of the classic definitions of the sulcus intermedius primus and secundus of Eberstaller (1884). While modern definitions retain the aipsJ label for the former—and credit it to the earlier definition by Jensen (1870)—the latter has been relabeled pips, which we identify in every hemisphere independent of the pAngs components (when present). For historical clarity, Bailey and colleagues (Bailey and von Bonin, 1951) reference that Eberstaller (1884) “borrowed” Jensen’s (Jensen, 1870) terminology: “Eberstaller (1884) who divided the inferior parietal lobule into three “arcs,” namely the supramarginal and angular gyri and the posterior parietal arc, recognized two intermediate sulci, “borrowing the term from Jensen, but understanding by it something quite different.”

This “borrowing” then led some subsequent authors to credit both Jensen and Eberstaller for the label. For example, Hrdlicka (1901) writes: “The supramarginal gyrus is fairly well defined on the left and is divided from the angular gyrus by a vertical branch proceeding from the interparietal sulcus (the sulcus intermedius primus, Jensen, Eberstaller).” pg. 478 And most recently, by ten Donkelaar and colleagues (2018) in which they write: “Clearly visible are the first and second intermediate parietal sulci of Jensen and Eberstaller (s.imdI and s.imdII, respectively).” These branches have also received additional labels, with some confusion relative to the caudal branches of the sts. For example, Bailey and colleagues (1951) identified three components of the aipsJ in which they write: “In brain *HI* the anterior part of the parallel sulcus shows several longer branches labeled simply 1-4. The posterior part breaks up into two rami, an anterior one (*pj*) and a posterior one (*ts*). The anterior branch connects by two subbranches (*pja* and *pjp*) with the intraparietal sulcus. The posterior branch anastomoses with *os*.”

Finally, recent work references shallow dimples between cSTS1 and cSTS2, which would be in the vicinity of our pAngs components. Specifically, Zlatkina and Petrides (2014) write: “In over a quarter of all examined hemispheres, a shallow sulcus or a set of dimples not connected with the IPS was observed between the first and second caudal branches of the superior temporal sulcus (27.5% of the left and 37.5% of the right hemispheres; Fig. 2a,d; electronic supplementary material, Fig. S1e)” pg. 4.

#### slocs/pAngs vs. F.I.P.r.int.1 and F.I.P.r.int.2

Recent work (Borne et al., 2020; Perrot et al., 2011) identified two intermediate rami of the IPS (F.I.P.r.int.1 and F.I.P.r.int.2) that were not defined in the present investigation. Crucially, the newly classified sulci here (slocs and pAngs) are distinguishable from the two F.I.P.r.int. in that the F.I.P.r.int. are branches coming off the main body of the IPS (Borne et al., 2020; Perrot et al., 2011), whereas the slocs/pAngs are predominantly non-intersecting (“free”) structures that never intersected with the IPS (**Supplementary Tables 1-4**).

## Supplementary Results

### Demographic variables were not included in the behavioral analysis

We did not include potentially relevant demographic measures of age, gender, and handedness as these did not reliably associate with our behavioral measures of interest (reasoning, age: *r* = −0.04, *p* = .74, gender: t = 1.01, *p* = .31, handedness: r = −0.003, *p* = .97; spatial orientation, age: *r* = 0.18, *p* = .14, gender: t = 1.54, *p* = .12, handedness: r = −0.05, *p* = 0.68; processing speed, age: *r* = −0.22, *p* = .06, gender: t = 0.07, *p* = .94, handedness: r = −0.09, *p* = .45).

### Hemispheric asymmetries in morphological, architectural, and functional features with regards to the slocs-v, cSTS3, and lTOS comparison

We observed a sulcus x metric x hemisphere interaction on the morphological and architectural features of the slocs-v (F(4.20, 289.81) = 4.16, η2 = 0.01, *p* = .002; the cSTS3 is discussed in the next section). Post hoc tests showed that this interaction was driven by the slocs-v being cortically thinner in the left than the right hemisphere (*p* < .001; **Fig. 3a**).

There was also a sulcus x network x hemisphere interaction on the functional connectivity profiles (using functional connectivity parcellations from (Kong et al., 2019) of the slocs-v and lTOS (F(32, 2144) = 3.99, η2 = 0.06, *p* < .001; the cSTS3 is discussed in the next section). Post hoc tests showed that this interaction was driven by three effects: (i) the slocs-v overlapped more with the Default C subnetwork in the left than the right hemisphere (*p* = .013), (ii) the lTOS overlapped more with Visual A subnetwork in the right than the left hemisphere (*p* = .002), and (iii) the lTOS overlapped more with the Visual B subnetwork in the left than the right hemisphere (*p* = .002; **Fig. 3b**).

### The caudal rami of the superior temporal sulcus are morphologically, architecturally, and functionally dissociable

As discussed in the historical section, though the three cSTS rami were most recently labeled by Segal and Petrides (2012), many neuroanatomists have labeled them throughout history in different species; **Fig. 1** and **Supplementary Fig. 5**). However, to our knowledge, it is not yet known whether these structures are distinguishable based on morphological, architectural, and functional features.

As described in the main text, we compared the morphological (depth and surface area) and architectural (gray matter thickness and myelination) features of these cSTS rami with an rm-ANOVA (within-participant factors: sulcus, metric, and hemisphere for standardized metric units). We observed a sulcus x metric interaction (F(3.48, 246.99) = 39.95, η2 = 0.36, *p* < .001). Post hoc tests showed that morphologically, the cSTS3 was deeper than the cSTS2 (*p* = .026) and cSTS1 (*p* < .001), while the cSTS2 and cSTS1 did not significantly differ (*p* = .12; **Supplementary Fig. 7a**). Further, the cSTS3 was smaller than the cSTS1 (*p* = .005) but not cSTS2 (*p* = .10), and cSTS1 and cSTS2 did not significantly differ (*p* = .99; **Supplementary Fig. 7a**). Architecturally, the cSTS3 was thinner than the cSTS2 and cSTS1 (*p*s < .001), but the cSTS2 and cSTS1 did not significantly differ (*p* = .74; **Supplementary Fig. 7a**). In addition, on myelination (i.e., the T1-w/T2-w ratio proxy), the cSTS3 was more myelinated than the cSTS2 and cSTS1 (*p*s < .001), but the cSTS2 and cSTS1 did not significantly differ (*p* = .11; **Supplementary Fig. 7a**).

It is also worth noting that there was a sulcus x metric x hemisphere interaction (F(4, 284.12) = 6.60, η2 = 0.08, *p* < .001). Post hoc tests showed that: (i) the cSTS3 was smaller (*p* < .001) and thinner (*p* = .025) in the left than the right hemisphere (**Supplementary Fig. 7a**), (ii) the cSTS2 was shallower (*p* = .004) and thicker (*p* < .001) in the right than left hemisphere (**Supplementary Fig. 7a**), and (iii) the cSTS1 was shallower (*p* < .001), smaller (*p* = .002), thinner (*p* = .001), and less myelinated (*p* < .001) in the left than the right hemisphere (**Supplementary Fig. 7a**).

Comparing the resting-state functional “connectivity fingerprints” (Kong et al., 2019) of the cSTS with an rm-ANOVA (within-participant factors: sulcus, network, and hemisphere for Dice coefficient overlap) revealed a sulcus x network interaction (F(32, 2208) = 88.31, η2 = 0.56, *p* < .001). Regarding subsequent post hoc test results, we only discuss effects related to networks that at least showed minor overlap with one cSTS (i.e., Dice ≥ .10). On the Auditory network, cSTS1 overlapped more than cSTS2 (*p* < .001; but not cSTS3: *p* = .57) and cSTS3 marginally more than cSTS2 (*p* = .052; **Supplementary Fig. 7a**). On the Control subnetworks, there was an superior-inferior difference: cSTS1 overlapped more with subnetworks B and C than both cSTS2 and cSTS3 (*p*s < .002), and cSTS2 overlapped more with subnetworks B and C than cSTS3 (*p*s < .006; **Supplementary Fig. 7b**). On the Default subnetworks, there was a different relationship for each subnetwork: i) cSTS2 overlapped more with Default subnetwork A than both cSTS1 and cSTS3 (*p*s < .001) and cSTS3 overlapped more than cSTS1 (*p* < .001), ii) cSTS1 and cSTS2 overlapped comparably with Default subnetwork B (*p* = .20), but both more than cSTS3 (*p*s < .001), and iii) cSTS3 overlapped more with Default subnetwork C than both cSTS1 and cSTS2 (*p*s < .001) and cSTS2 overlapped more than cSTS1 (*p* < .001; **Supplementary Fig. 7b**). On the Dorsal Attention A subnetwork, cSTS3 overlapped more than cSTS1 and cSTS2 (*p*s < .001) and cSTS2 overlapped more than cSTS1 (*p* = .001; **Supplementary Fig. 7b**). On the Temporal-Parietal Network, cSTS1 overlapped more than cSTS2 and cSTS3 (*p*s < .001), and cSTS2 and cSTS3 did not differ significantly: *p* = .25; **Supplementary Fig. 7b**). On the Ventral Attention B subnetwork, cSTS1 overlapped more than cSTS2 (*p* < .001) and marginally more than cSTS3 (*p* = .064), and cSTS2 and cSTS3 did not differ significantly (*p* = .11; **Supplementary Fig. 7b**). Finally, on the Visual A subnetwork, cSTS3 overlapped more than both cSTS1 and cSTS2 (*p*s < .001), and cSTS1 and cSTS2 did not significantly differ (*p* = .15; **Supplementary Fig. 7b**).

There was also a sulcus x network x hemisphere interaction (F(32, 2208) = 12.26, η2 = 0.15, *p* < .001). Post hoc tests showed differences for each cSTS component. Here, the cSTS1 overlapped more with the Auditory network (*p* < .001), less with the Control B subnetwork (*p* < .001), more with the Control C subnetwork (*p* < .001), less with the Default B subnetwork (*p* < .001), more with the Default C subnetwork (*p* < .001), more with the Ventral Attention B subnetwork (*p* < .001), and more with the Visual A subnetwork (*p* = .024) in the right than in the left hemisphere (**Supplementary Fig. 7b**). In addition, the cSTS2 overlapped more with the Control B subnetwork (*p* < .001), more with the Control C subnetwork (*p* < .001), less with the Default B subnetwork (*p* < .001), and less with the Temporal-Parietal network (*p* = .011) in the right than in the left hemisphere (**Supplementary Fig. 7b**). Finally, the cSTS3 overlapped more with the Control B subnetwork (*p* = .002), less with the Default B subnetwork (*p* = .014), more with the Default C subnetwork (*p* = .022), less with the Ventral Attention B subnetwork (*p* = .029) in the right than in the left hemisphere (**Supplementary Fig. 7b**).

Altogether, these data indicate that the cSTS3 is moderately morphologically (deeper and smaller) and largely architecturally (thinner and more myelinated) distinguishable from the more dorsal cSTS (cSTS1 and cSTS2), which largely do not differ in these metrics. In addition, the three cSTS all differ in their relationship to resting-state functional connectivity networks.

Specifically, the cSTS1 overlaps with Auditory, Control (B and C), Default (B), Temporal-Parietal, Ventral Attention (B) networks/subnetworks, the cSTS2 overlaps with Default (A and B) subnetworks, and the cSTS3 overlaps with Default (C), Dorsal Attention (A), and Visual (A) subnetworks. Regarding the cSTS3-related results, these findings especially support the notion that the cSTS3 is an anatomical and functional transition region between the lateral parietal and lateral occipital cortices (Glasser et al., 2016); **Fig. 3**).

### The anterior intermediate parietal sulcus of Jensen and posterior intermediate parietal sulcus are morphologically, architecturally, and functionally dissociable

As also discussed in the historical section, there are two intermediate parietal sulci (ips) in LPC: the anterior ips of Jensen (aipsJ; Bailey and von Bonin, 1951; Eberstaller, 1884; Jensen, 1870; von Economo and Koskinas, 1925; Zlatkina and Petrides, 2014) and the posterior ips (pips; Petrides, 2019; Zlatkina and Petrides, 2014). Further, to our knowledge and as with the three cSTS, it is not known whether the two ips are distinguishable based on morphological, architectural, and functional features.

Comparing the morphological (depth and surface area) and architectural (gray matter thickness and myelination) features of the ips with an rm-ANOVA (within-participant factors: sulcus, metric, and hemisphere for standardized metric units) revealed a sulcus x metric interaction (F(1.58, 112.15) = 93.00, η2 = 0.57, *p* < .001; no sulcus x metric x hemisphere interaction: *p* = .76). Post hoc tests showed that, morphologically, the aipsJ was shallower than the pips (*p* < .001), but the two ips were comparably sized (*p* = .58; **Supplementary Fig. 7c**). Further, these post hoc tests showed that, architecturally, the aipsJ was thicker and less myelinated than the pips (*p*s < .001; **Supplementary Fig. 7c**).

In addition, comparing the resting-state functional “connectivity fingerprints” (Kong et al., 2019) of the ips with an rm-ANOVA (within-participant factors: sulcus, network, and hemisphere for Dice coefficient overlap) revealed a sulcus x network interaction (F(16, 1104) = 61.73, η2 = 0.47, *p* < .001). Post hoc tests showed that: (i) the pips overlapped more with the Control A subnetwork (*p* < .001), (ii) the aipsJ overlapped more with the Control B and C subnetworks (*p* < .001), (iii) the aipsJ overlapped more with the Default A and B subnetworks (*p*s < .001), (iii) the pips overlapped more with the Dorsal Attention A subnetwork (*p* < .001), and (iv) the pips overlapped more with the Visual A subnetwork (*p* < .001; **Supplementary Fig. 7d**).

There was also a sulcus x network x hemisphere interaction (F(16, 1104) = 6.70, η2 = 0.09, *p* < .001). Post hoc tests showed differences for both ips. First, the aipsJ overlapped more with the Control A subnetwork (*p* = .011), less with the Default A subnetwork (*p* = .001), less with the Default B subnetwork (*p* = .041), and more with the Dorsal Attention A subnetwork (*p* = .033) in the right than the left hemisphere (**Supplementary Fig. 7d**). Second, the pips overlapped less with the Control A subnetwork (*p* = .003) and more with the Dorsal Attention A subnetwork (*p* = .011) in the right than the left hemisphere (**Supplementary Fig. 7d**).

Altogether, these data indicate that the two ips are morphologically, architecturally, and functionally dissociable structures. The aipsJ is smaller, thicker, and less myelinated than the pips. The aipsJ also overlaps more with Control (B and C) and Default (A and B) subnetworks, whereas the pips overlaps more with a single Control (A), Dorsal Attention (A), and Visual (A) subnetworks.

## Data availability

The processed data required to perform all statistical analyses and reproduce all Figures, as well as the probability maps, are available on GitHub (https://github.com/cnl-berkeley/stable_projects). Anonymized HCP neuroimaging data are publicly available on ConnectomeDB (db.humanconnectome.org). Raw data will be made available from the corresponding author upon request.

## Code availability

The code is available on GitHub (https://github.com/cnl-berkeley/stable_projects) and Open Science Framework (https://osf.io/7fwqk/).

## Acknowledgments

This research was supported by NSF CAREER Award 2042251 (PI Weiner) and NIH MSTP Grant T32 GM140935 (Willbrand). Neuroimaging and behavioral data were provided by the HCP, WU-Minn Consortium (PIs: David Van Essen and Kamil Ugurbil; NIH Grant 1U54-MH-091657) funded by the 16 NIH Institutes and Centers that support the NIH Blueprint for Neuroscience Research, and the McDonnell Center for Systems Neuroscience at Washington University. We thank Jacob Miller, Benjamin Parker, and Willa Voorhies for helping develop the analysis pipelines implemented in this project. We also thank the HCP researchers for participant recruitment and data collection and sharing, as well as the participants who participated in the study.

## Supplementary Tables

**Supplementary Table 1.**
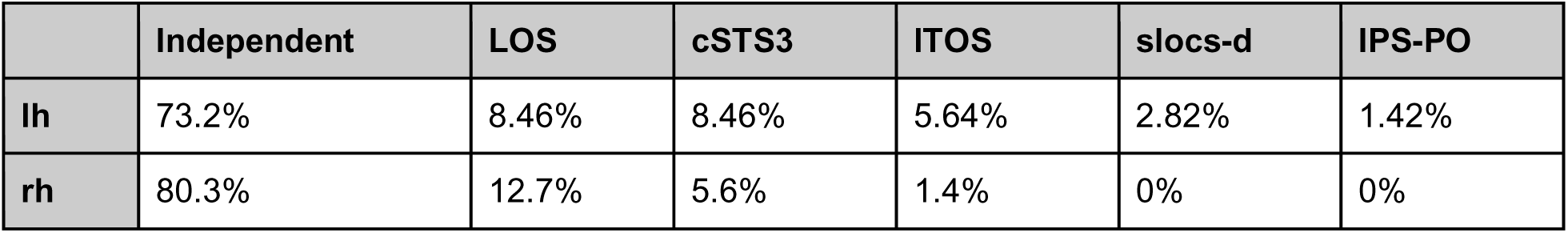
Slocs-v sulcal types. This table displays the incidence rates of how often the slocs-v intersects with a surrounding sulcus (as a percentage; out of 71 for both hemispheres). Independent means there are no intersections. These rates are highly similar between hemispheres (r = .99, *p* < .0001). The LOS (lateral occipital sulcus) is not described in the main text but is a sulcus ventral to lTOS, cSTS3, and slocs-v in lateral occipital cortex (Petrides, 2019).

**Supplementary Table 2.** Slocs-d sulcal types. This table displays the incidence rates of how often the slocs-d intersects with a surrounding sulcus (as a percentage; out of 50 in the left hemisphere and 48 in the right hemisphere). Independent means there are no intersections. These rates are highly similar between hemispheres (r = .99, *p* < .0001).

|  | Independent | cSTS2 | cSTS3 | slocs-v | IPS-PO | pAngs-v | pips |
| --- | --- | --- | --- | --- | --- | --- | --- |
| lh | 80% | 6% | 4% | 4% | 2% | 2% | 2% |
| rh | 75.5% | 2.05% | 4.1% | 0% | 8.15% | 6.1% | 4.1% |

**Supplementary Table 3.**
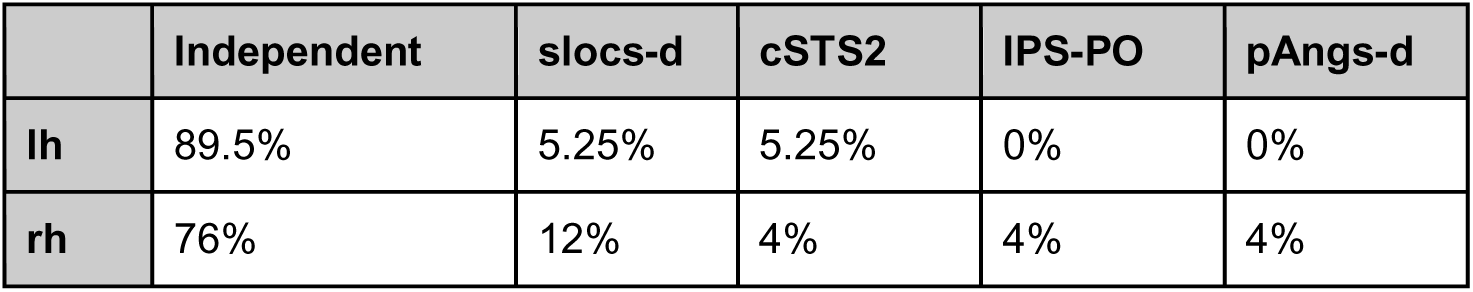
pAngs-v sulcal types. This table displays the incidence rates of how often the pAngs-v intersects with a surrounding sulcus (as a percentage; out of 19 in the left hemisphere and 26 in the right hemisphere). Independent means there are no intersections. These rates are highly similar between hemispheres (r = .99, *p* = .0003).

|  | Independent | slocs-d | cSTS2 | IPS-PO | pAngs-d |
| --- | --- | --- | --- | --- | --- |
| lh | 89.5% | 5.25% | 5.25% | 0% | 0% |
| rh | 76% | 12% | 4% | 4% | 4% |

**Supplementary Table 4.**
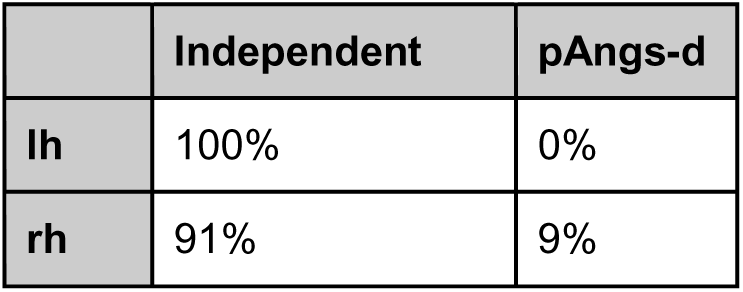
pAngs-d sulcal types. This table displays the incidence rates of how often the pAngs-d intersects with a surrounding sulcus (as a percentage; out of 8 in the left hemisphere and 11 in the right hemisphere). Independent means there are no intersections.

**Supplementary Table 5.**
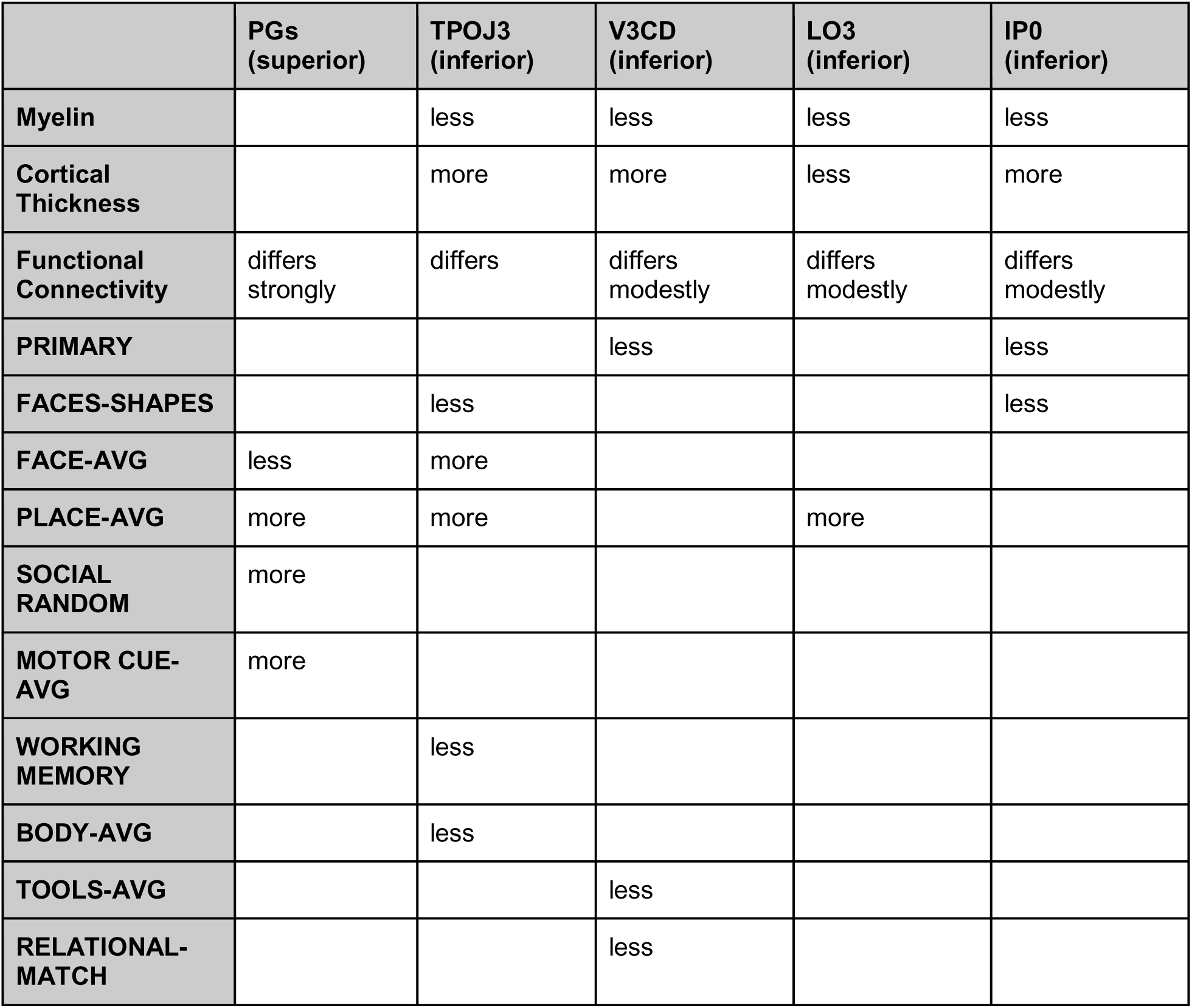
Differences between HCP-MMP area PGp—the area that the slocs-v co-localized with at the group and probabilistic level—and surrounding areas. This table displays the values of PGp relative to each of the surrounding regions (details are from the Supplementary Neuroanatomical Results section in Glasser et al., 2016). For example, if PGp is less myelinated than a region that box will say “less.” The location of each region relative to PGp is in parenthesis (superior indicates the region is above PGp, etc.). If an area is blank that difference was not stated in (Glasser et al., 2016). Functional contrasts are fully capitalized.

|  | **PGs**<br>**(superior)** | **TPOJ3**<br>**(inferior)** | **V3CD**<br>**(inferior)** | **LO3**<br>**(inferior)** | **IP0**<br>**(inferior)** |
| --- | --- | --- | --- | --- | --- |
| **Myelin** |  | less | less | less | less |
| **Cortical**<br>**Thickness** |  | more | more | less | more |
| **Functional**<br>**Connectivity** | differs<br>strongly | differs | differs<br>modestly | differs<br>modestly | differs<br>modestly |
| **PRIMARY** |  |  | less |  | less |
| **FACES-SHAPES** |  | less |  |  | less |
| **FACE-AVG** | less | more |  |  |  |
| **PLACE-AVG** | more | more |  | more |  |
| **SOCIAL**<br>**RANDOM** | more |  |  |  |  |
| **MOTOR CUE-**<br>**AVG** | more |  |  |  |  |
| **WORKING**<br>**MEMORY** |  | less |  |  |  |
| **BODY-AVG** |  | less |  |  |  |
| **TOOLS-AVG** |  |  | less |  |  |
| **RELATIONAL-**<br>**MATCH** |  |  | less |  |  |

## Supplementary Figures

**Supplementary Fig. 1.**
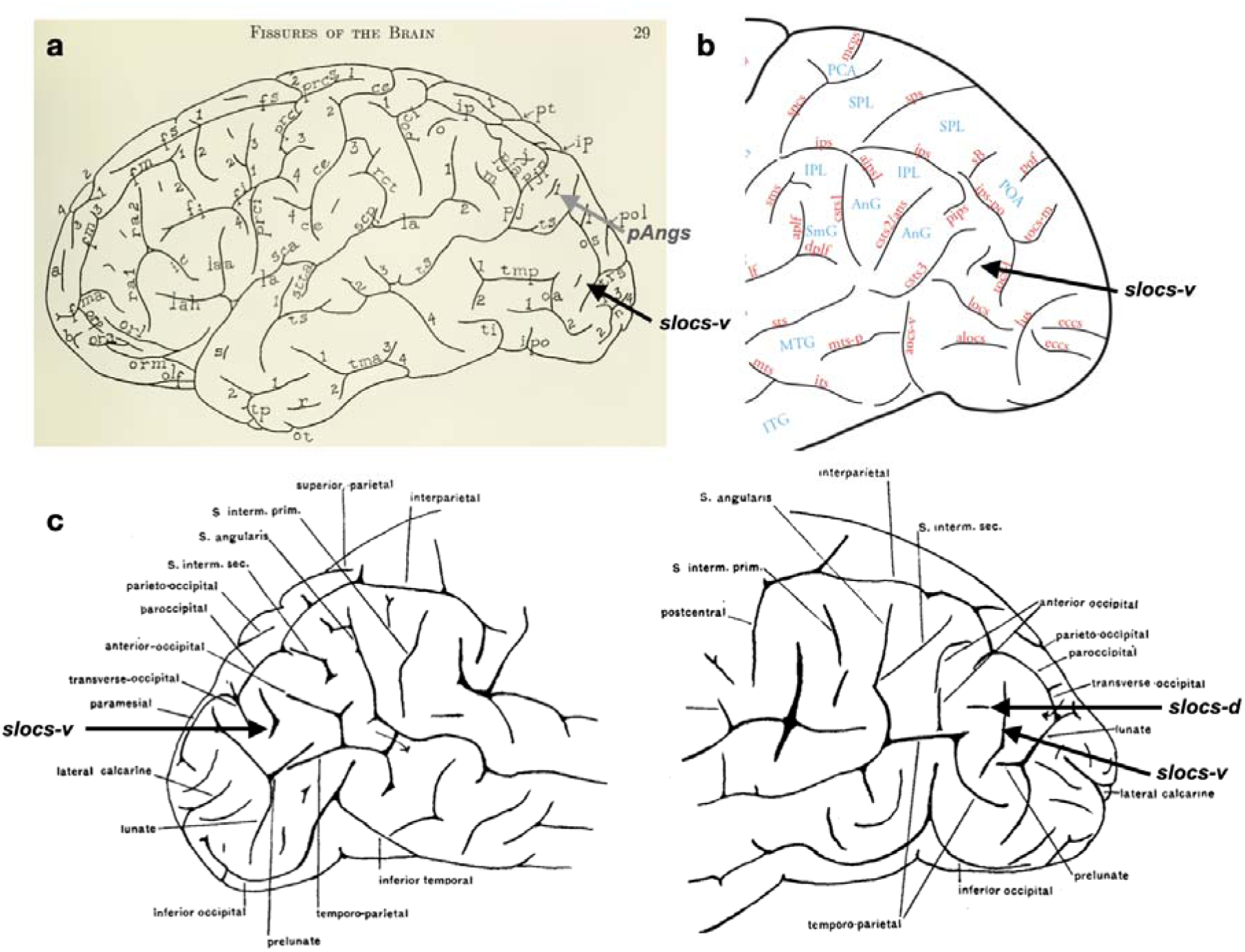
Supralateral occipital (slocs) and posterior angular (pAng) sulci relative to classic and modern sulcal definitions—References 1. **a.** Sulcal definitions from Bailey et al. (1951). The black arrow indicates a depicted, but unlabeled sulcus in the vicinity of our slocs-v. The gray arrow indicates a sulcus labeled “1” in the vicinity of our pAngs components. As Bonin et al. (1951) write: “Short, isolated dimples and sulci are given letters from a to z.” Numbers were given to rami. Note that instead of identifying the three branches of the STS, they identify additional anterior (pja) and posterior (pjp) branches of the aipsJ (what they refer to as *pj*). **b.** A depicted, but unlabeled slocs-v from the most recent atlas to include tertiary sulci from Petrides (2019). **c.** Depicted, but unlabeled slocs-v and slocs-d from Connolly (1950).

**Supplementary Fig. 2.**
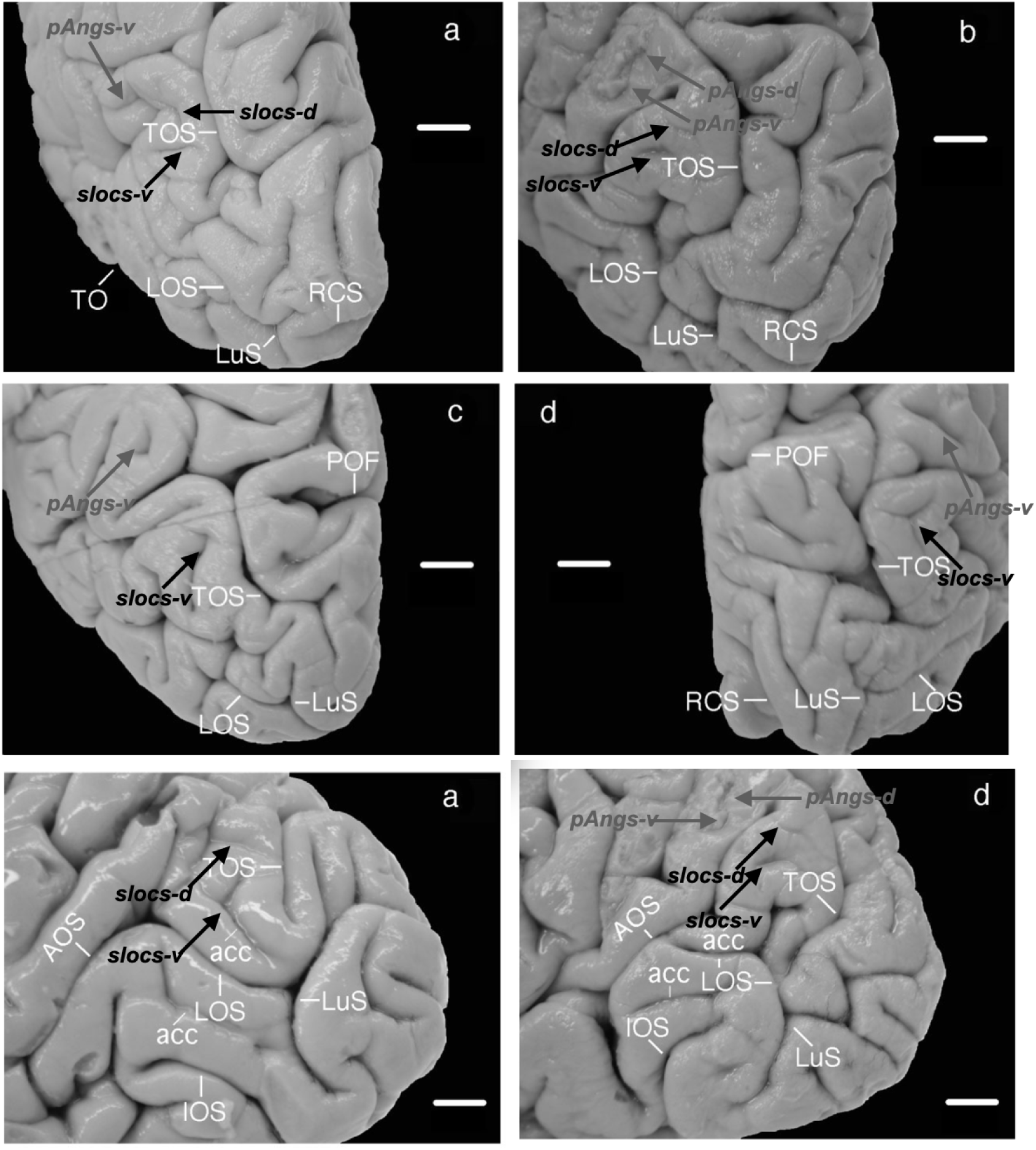
Supralateral occipital (slocs) and posterior angular (pAng) sulci relative to classic and modern sulcal definitions—Reference 2. Example postmortem hemispheres from Iaria and Petrides (2007) depicting the unlabeled slocs (black arrows) and pAngs (gray arrows) components.

**Supplementary Fig. 3.**
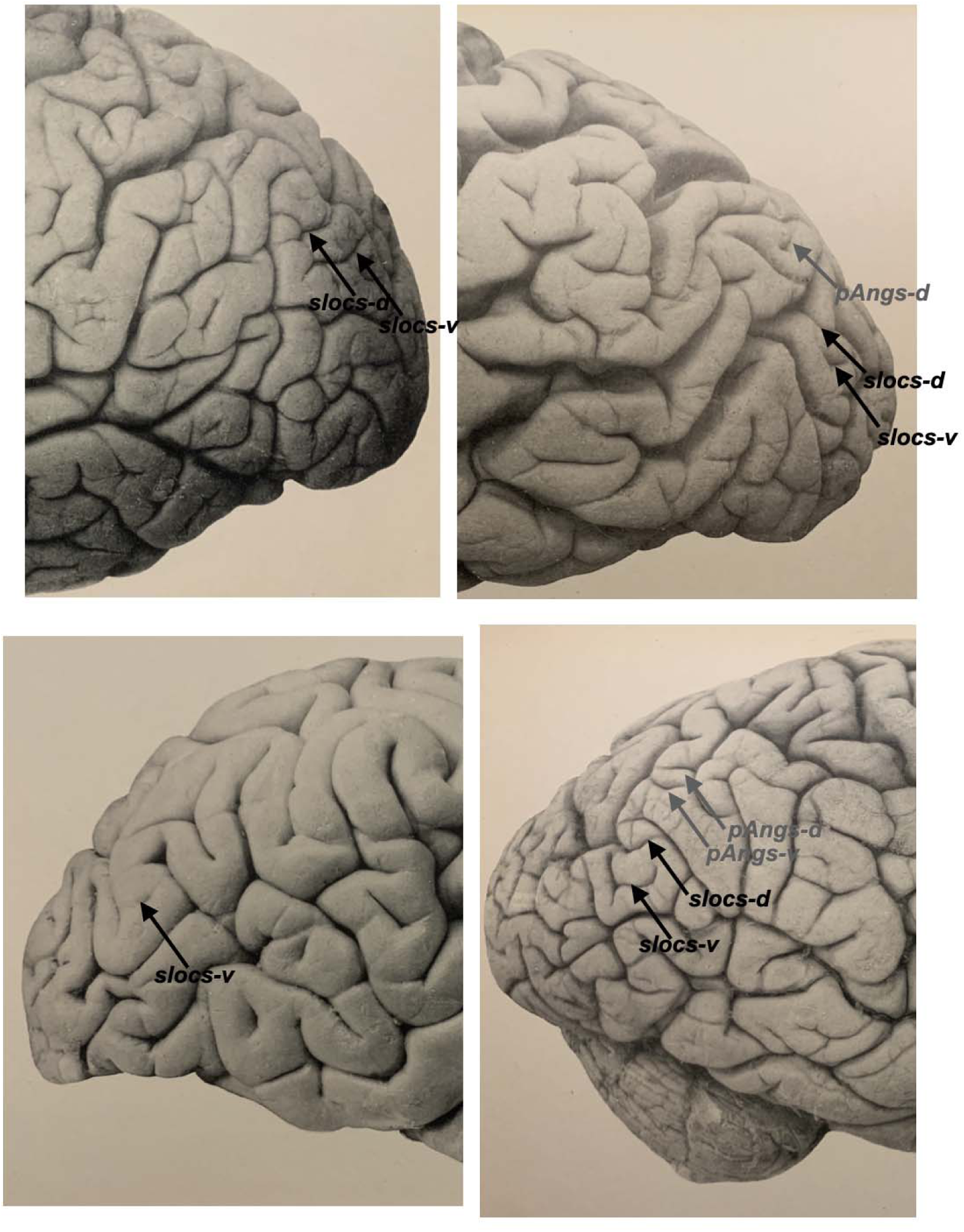
Supralateral occipital (slocs) and posterior angular (pAng) sulci relative to classic and modern sulcal definitions—Reference 3. Four example postmortem hemispheres from Retzius (1896) depicting the slocs (black arrows) and pAngs (gray arrows).

**Supplementary Fig. 4.**
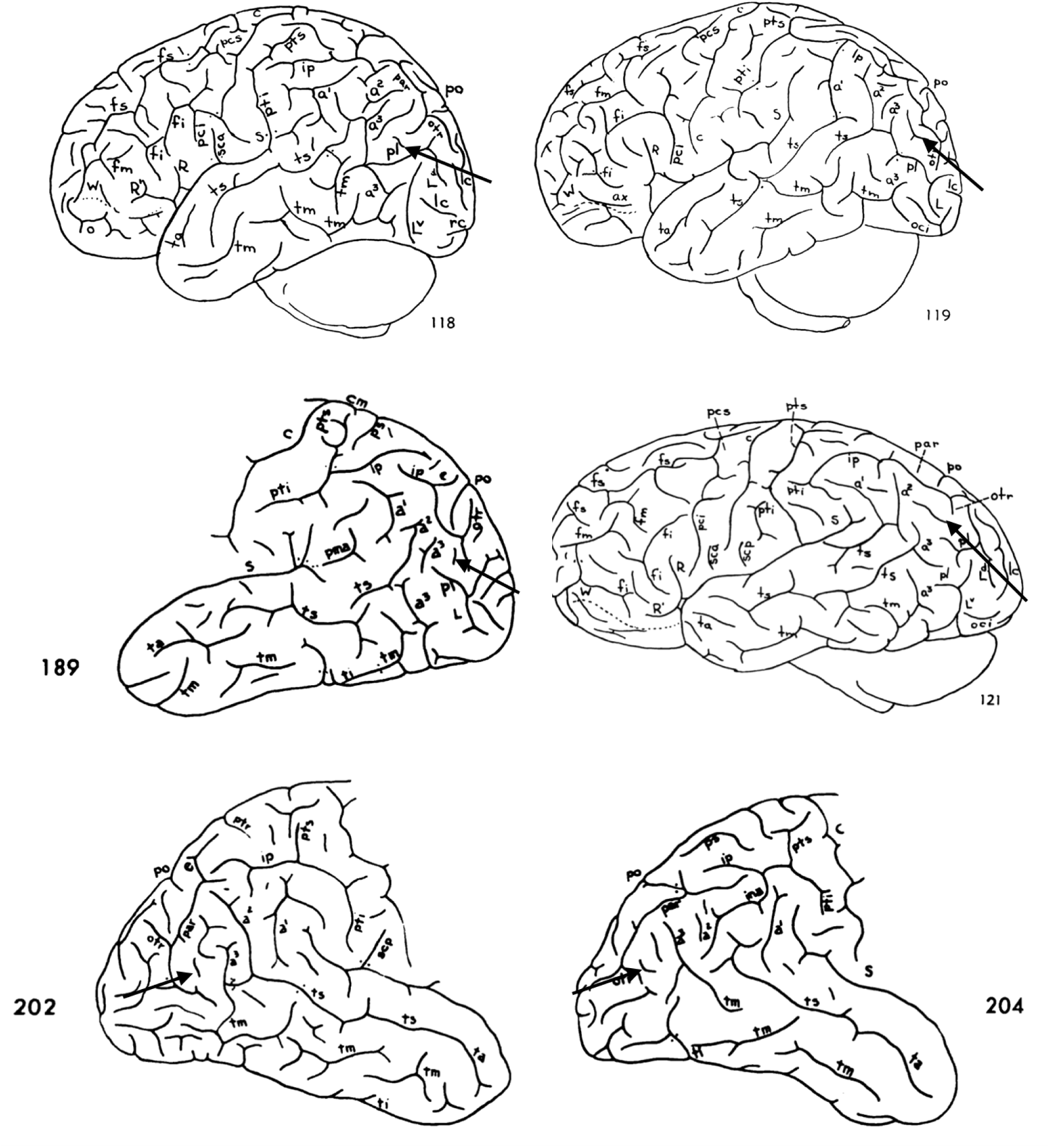
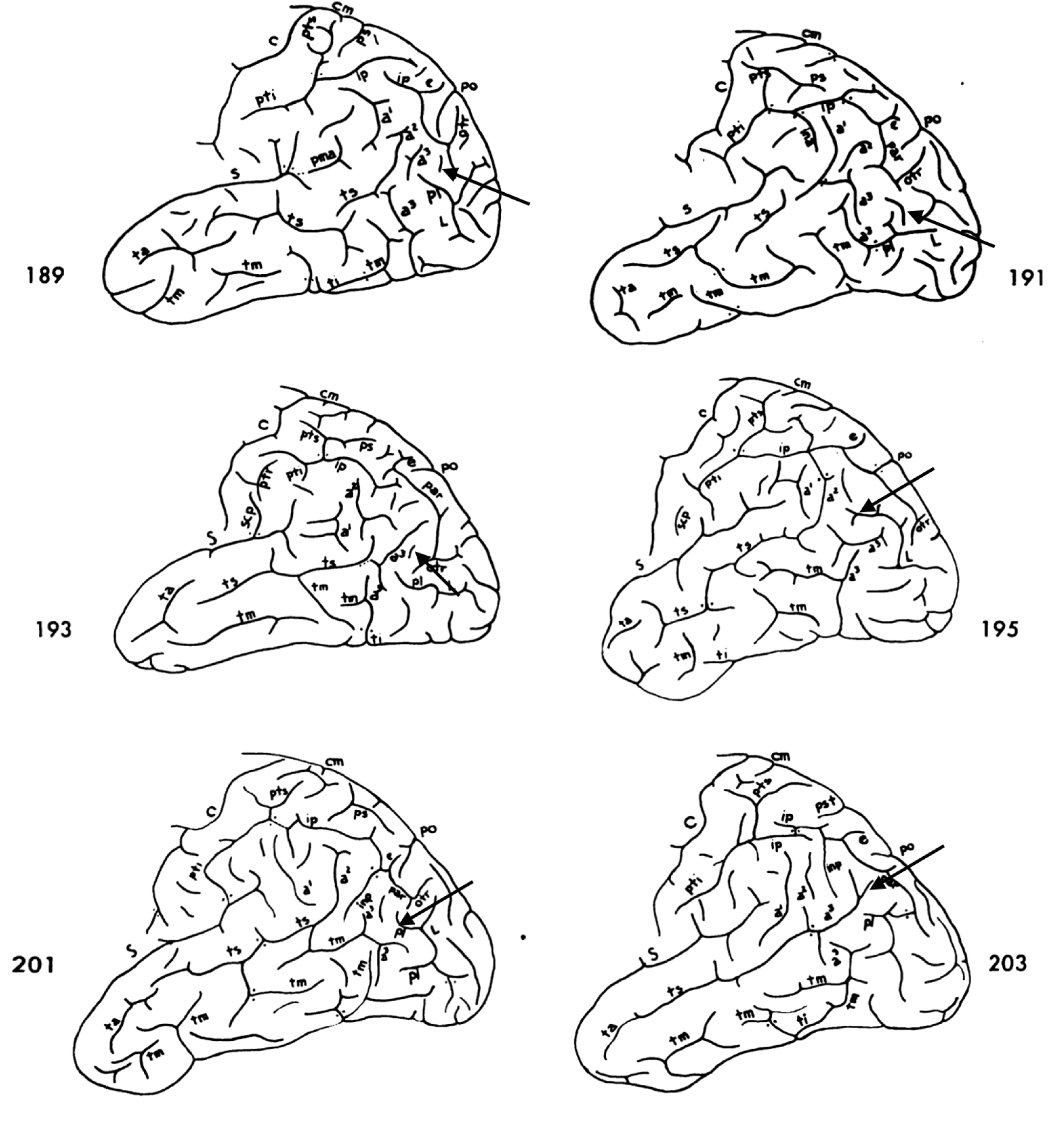
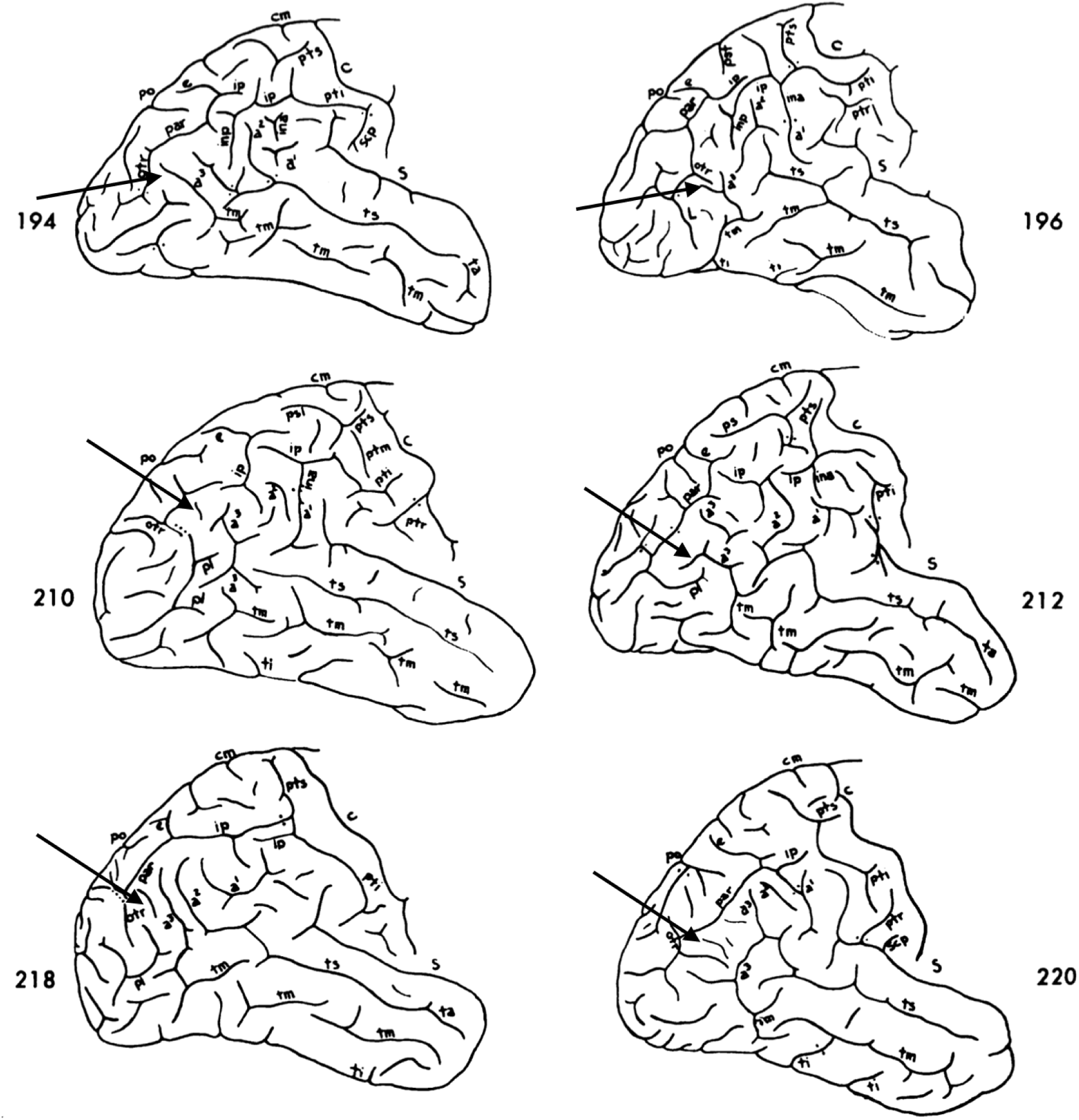
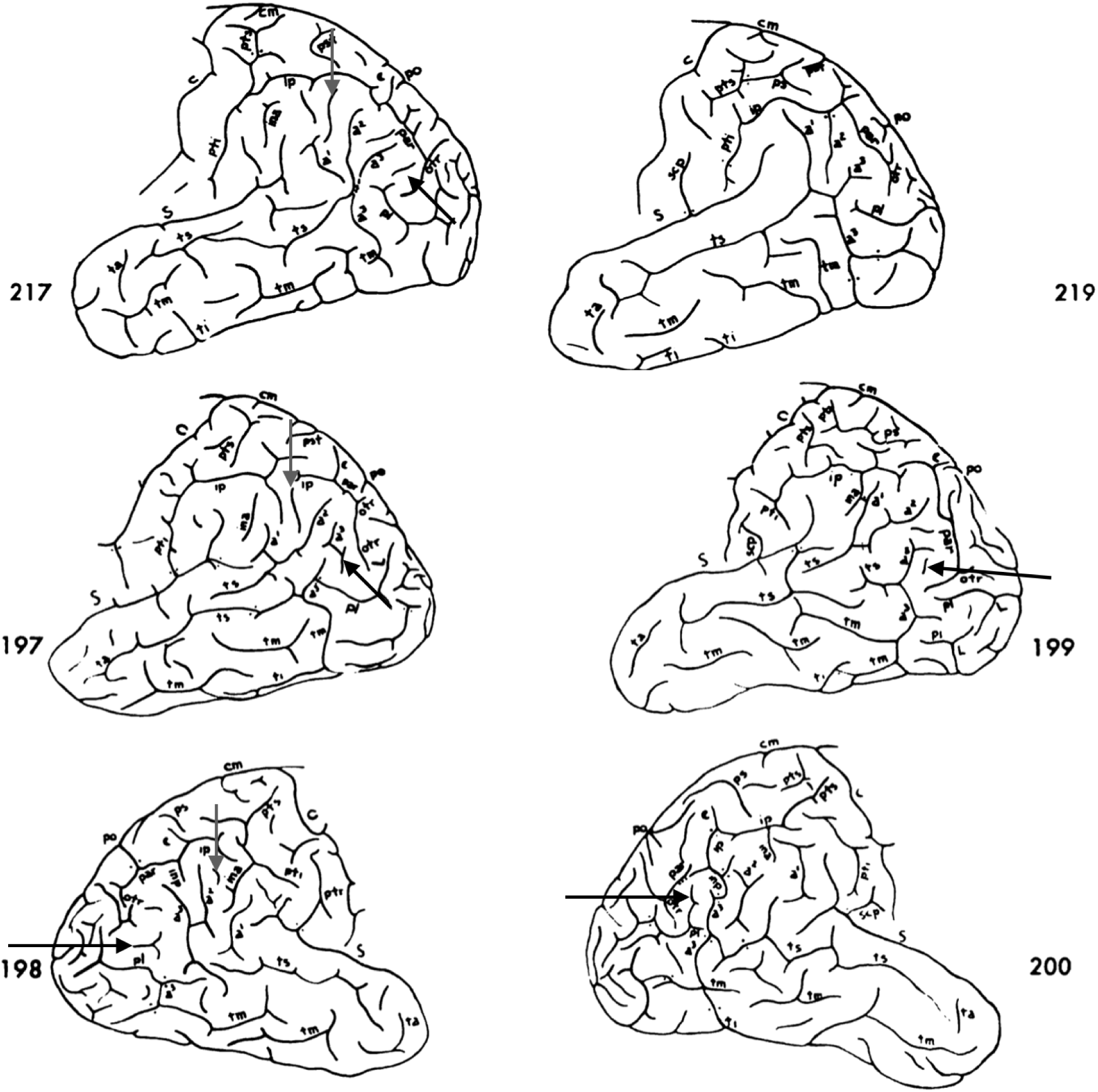
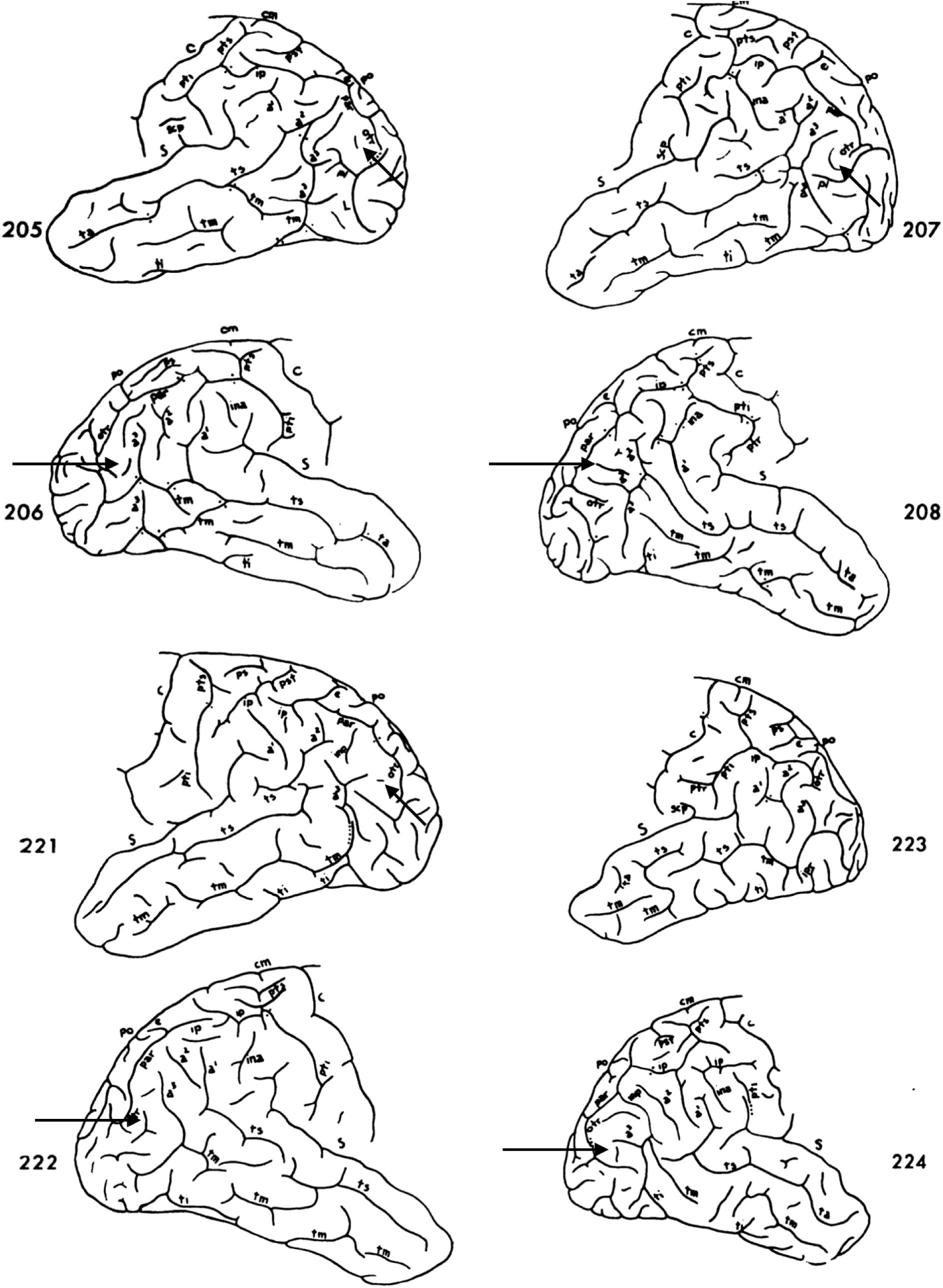
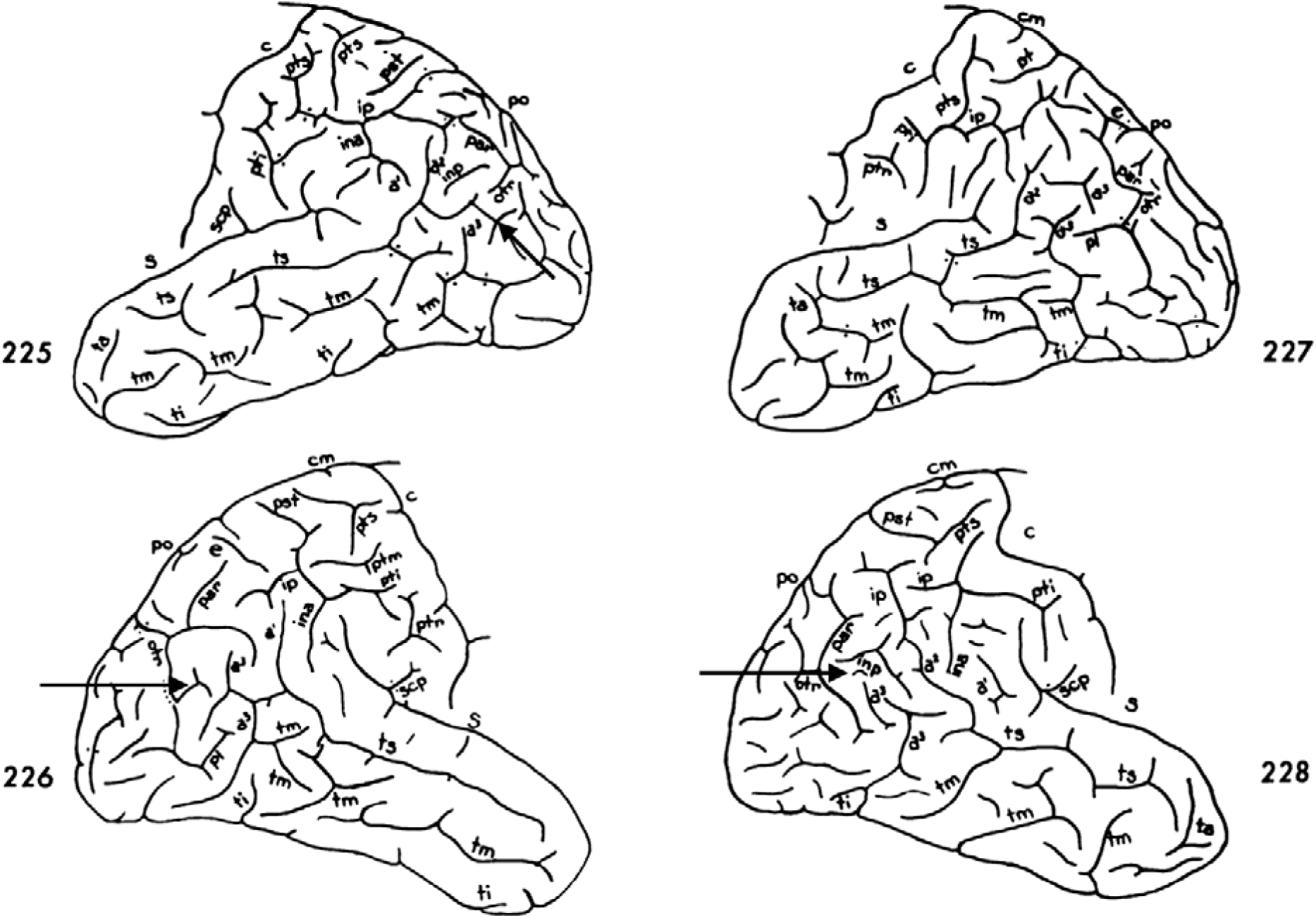
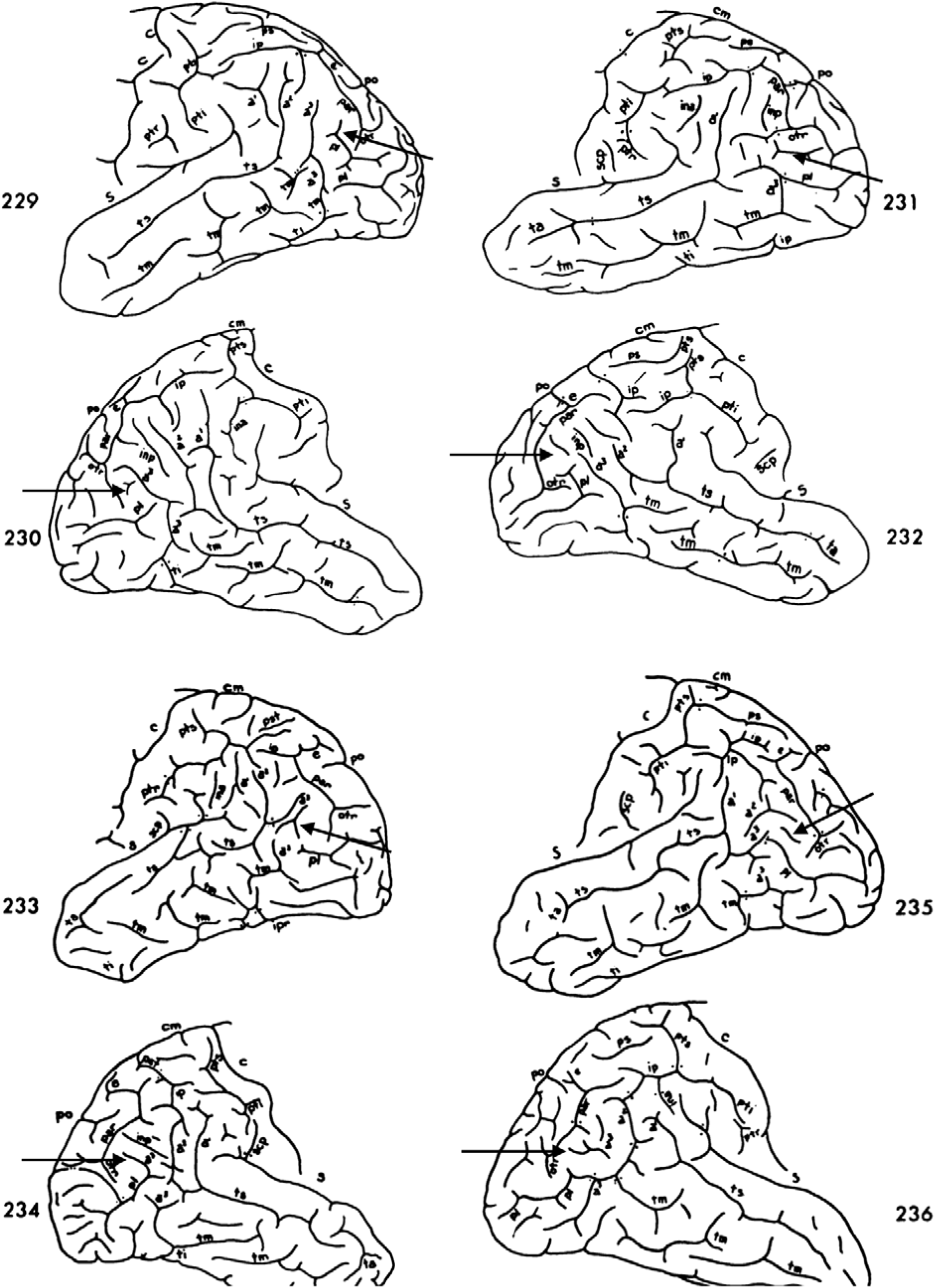
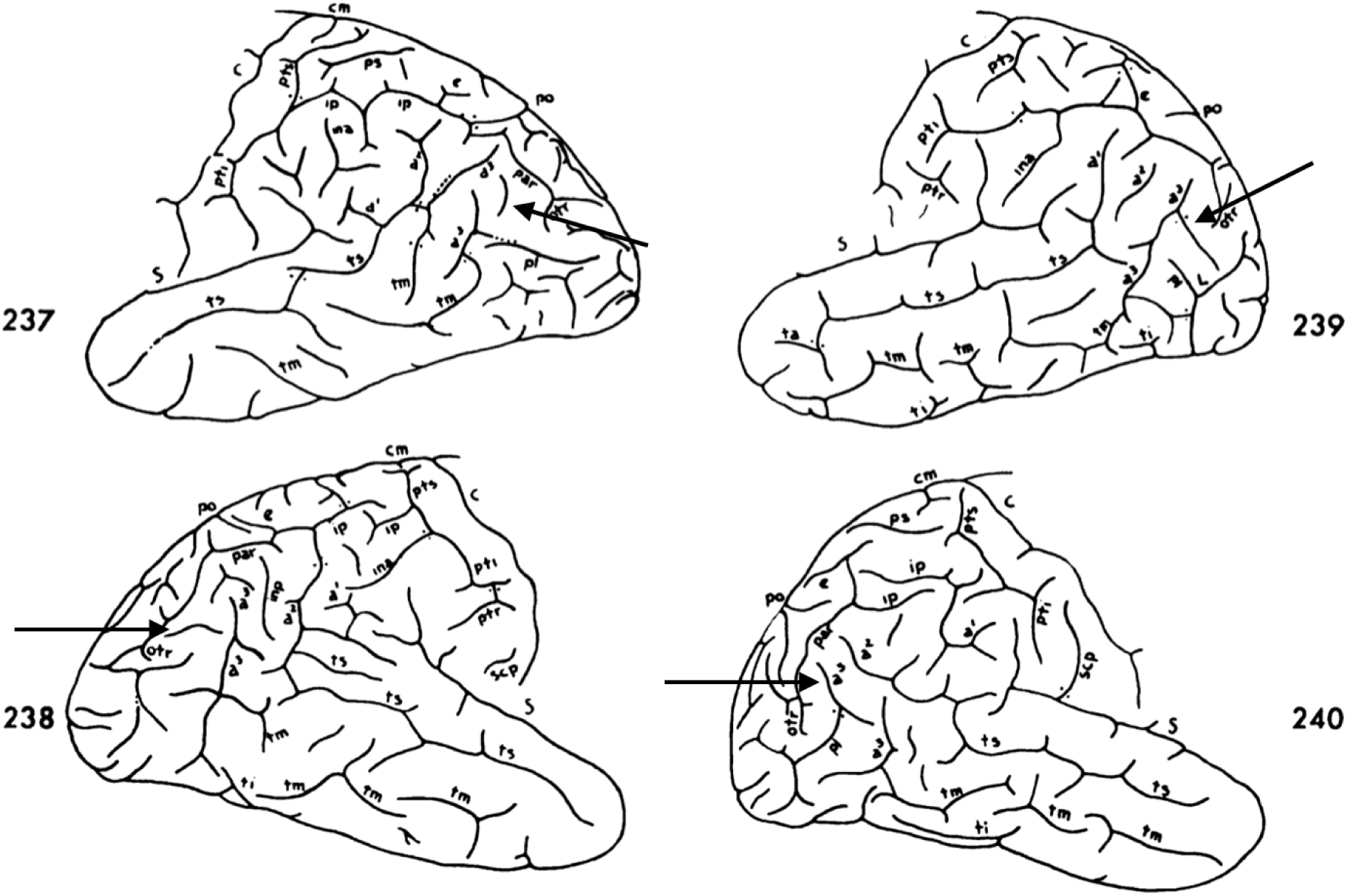
Ventral supralateral occipital sulcus (slocs-v) in human hemispheres from Connolly (1950). 50 example hemispheres from Connolly (1950) depicting an unlabeled slocs-v (black arrow) 94% of the time (47/50).

**Supplementary Fig. 5.**
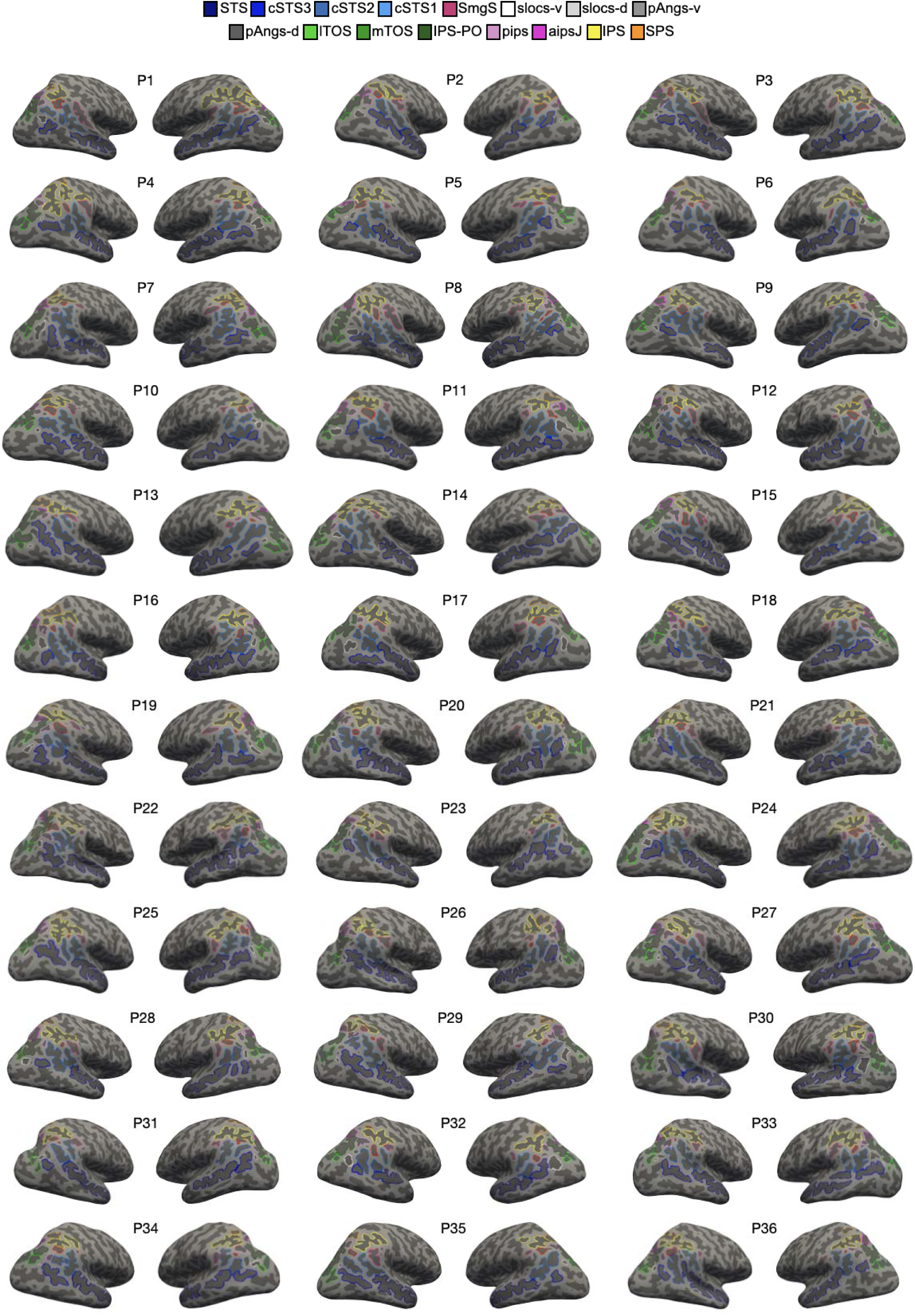
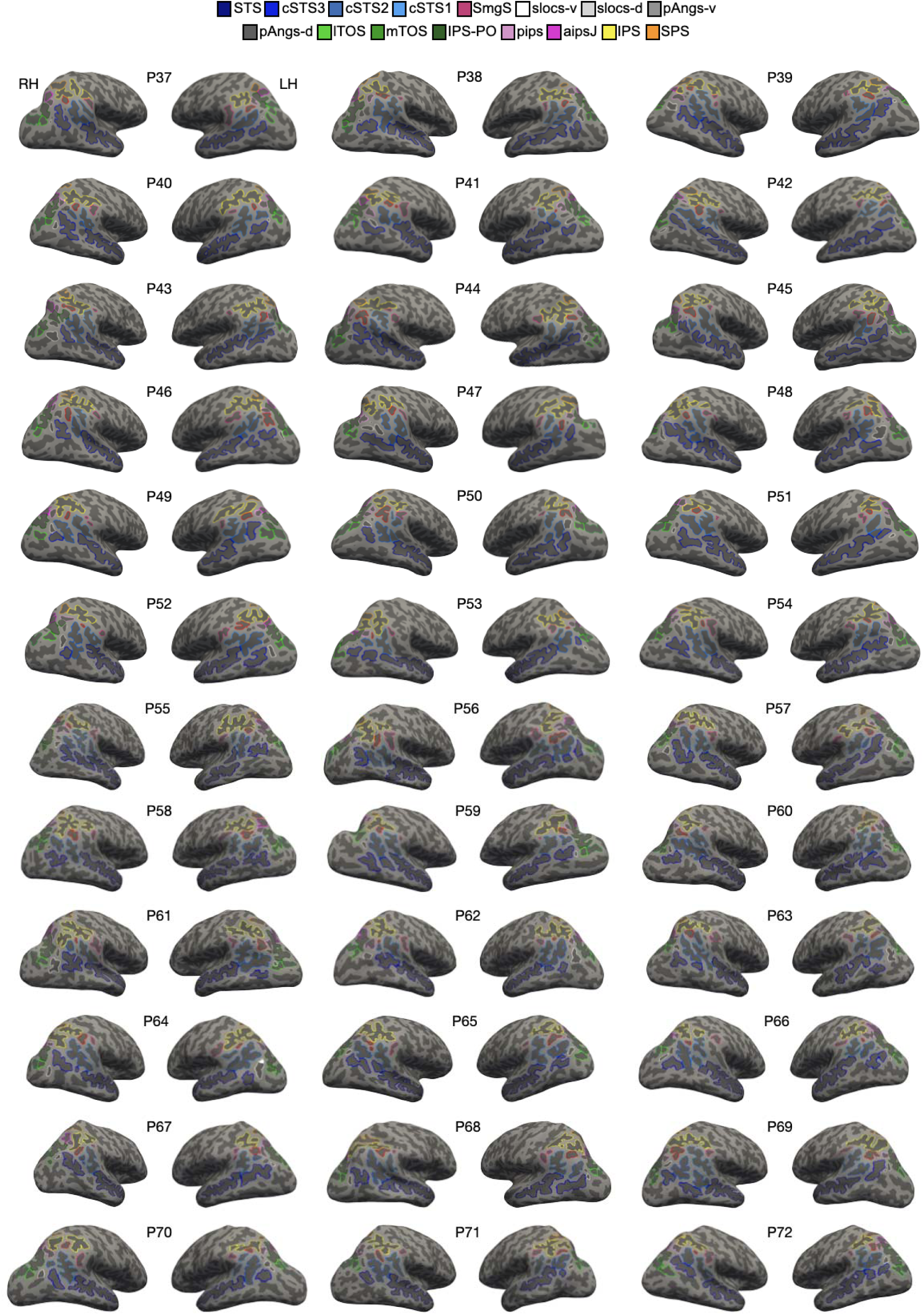
All 2176 LPC/LPOJ sulcal definitions across 72 participants (144 hemispheres). Each sulcus is displayed on the left (LH, right surfaces) and right (RH, left surfaces) inflated cortical surfaces for each participant (P) in FreeSurfer 6.0.0, with the label displayed as an outline according to the key at the top.

**Supplementary Fig. 6.**
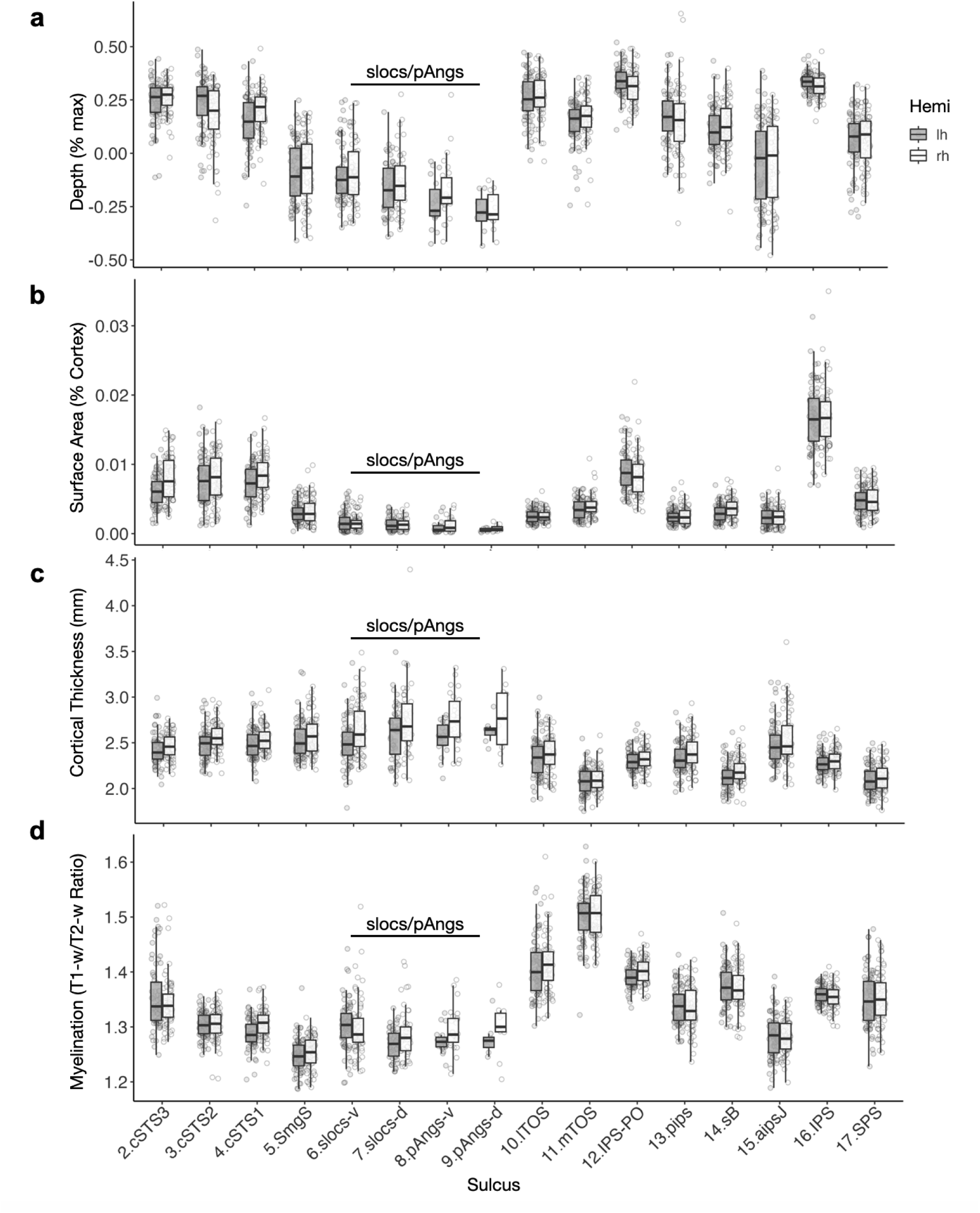
The slocs and pAngs ventral and dorsal components are among the smallest and shallowest structures in LPC/LPOJ. **a.** Box plots displaying depth (% maximum cortical depth) as a function of sulcus (x-axis) and hemisphere [left hemisphere (lh; black) and right hemisphere (rh; white)]. Individual dots represent values for individual participants. The newly-identified slocs and pAngs ventral and dorsal components are identified with the horizontal black line. We did not include STS in these plots given that it primarily resides outside the cortical expanse of interest (i.e., LPC/LPOJ). **b.** Same as a, but for surface area (normalized to % cortex surface area). **c.** Same as a, except for cortical thickness (mm). **d.** Same as a, except for myelination (T1w/T2w ratio).

**Supplementary Fig. 7.**
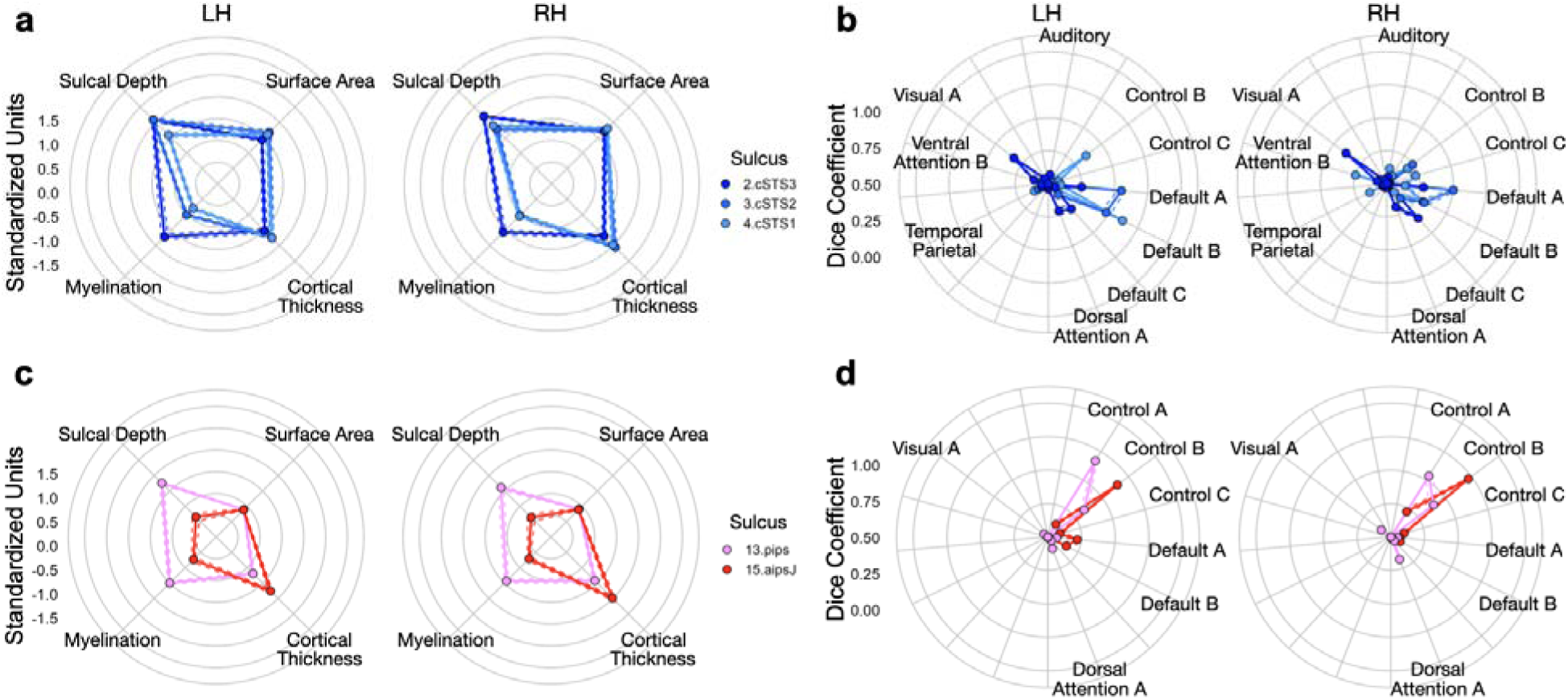
The three caudal rami of the superior temporal sulcus and intermediate parietal sulci are dissociable neuroanatomical structures. **a.** Radial plot displaying the morphological (upper metrics: depth, surface area) and architectural (lower metrics: cortical thickness, myelination) features of the caudal rami of the superior temporal sulcus (cSTS1 to 3, light to dark blue). Each dot and solid line represents the mean. The dashed lines indicate ± standard error. These features are colored b sulcus (see key). Metrics are standardized in order to be visualized on the same axis. **b.** Radial plot displaying the connectivity fingerprints of these three sulci: the Dice Coefficient overlap (values from 0-1) between each component and individual-level functional connectivity parcellations (Kong et al., 2019). The networks that each sulcus overlaps with (Dice > .10 for at least one sulcus) and present inter-sulcal differences are shown. **c.** Same as a, except for the anterior intermediate parietal sulcus of Jensen (aipsJ; red) and posterior intermediate parietal sulcus (pips; pink). **d.** *S*ame as b, except for the aipsJ and pips.

## Footnotes

1* Smith (1907) writes: “The whole of the area between the sulcus occipitalis lateralis (i.e. praelunatus) and the sulcus occipitalis inferior is often occupied by a cortical area indistinguishable from and continuous with the area peristria-ta; but part of this region (marked “ AR. TEM. occ.” in fig. 2) occasionally exhibits a faint doubling of the line of Baillarger, which calls for its separation from that area.” pg. 243

